# HIV-1 binds dynein directly to hijack microtubule transport machinery

**DOI:** 10.1101/2023.08.29.555335

**Authors:** Somayesadat Badieyan, Drew Lichon, Michael P. Andreas, John P. Gillies, Wang Peng, Jiong Shi, Morgan E. DeSantis, Christopher R. Aiken, Till Böcking, Tobias W. Giessen, Edward M. Campbell, Michael A. Cianfrocco

## Abstract

Viruses exploit host cytoskeletal elements and motor proteins for trafficking through the dense cytoplasm. Yet the molecular mechanism that describes how viruses connect to the motor machinery is unknown. Here, we demonstrate the first example of viral microtubule trafficking from purified components: HIV-1 hijacking microtubule transport machinery. We discover that HIV-1 directly binds to the retrograde microtubule-associated motor, dynein, and not via a cargo adaptor, as previously suggested. Moreover, we show that HIV-1 motility is supported by multiple, diverse dynein cargo adaptors as HIV-1 binds to dynein light and intermediate chains on dynein’s tail. Further, we demonstrate that multiple dynein motors tethered to rigid cargoes, like HIV-1 capsids, display reduced motility, distinct from the behavior of multiple motors on membranous cargoes. Our results introduce a new model of viral trafficking wherein a pathogen opportunistically ‘hijacks’ the microtubule transport machinery for motility, enabling multiple transport pathways through the host cytoplasm.

## Introduction

Due to the crowded nature and large size of eukaryotic cells, diverse processes, including organelle trafficking, mRNA localization, and cell division, take advantage of microtubule-based trafficking systems^1^. The directional cargo transport along microtubules is accomplished by the actions of opposite-directed motor proteins kinesin^2^ and dynein^3^.

The size of viral particles and the high density of the cytoplasm force viral pathogens to evolve mechanisms for hijacking and manipulating host transport machinery to navigate the host cell and support their replication^1,4^. One example is the human immunodeficiency virus type 1 (HIV-1)^5^. Nearly 20 years ago, live-cell imaging^6,7^ showed that HIV-1 exploits the microtubule network to move throughout the cell. Kinesin-1 or dynein motor protein depletion inhibits HIV-1 trafficking in the cytoplasm, nuclear translocation, uncoating, and infection^8,9^, showing that microtubule-associated motors play an integral part in the HIV-1 life cycle.

Despite visualizing the connection between HIV-1 and microtubules in cells, the molecular basis for the interaction remains poorly defined. For instance, a suite of perturbations show that dynein is required for motility, but fail to clearly point to a mechanism of attachment: 1) microinjection of an inhibitory antibody against the dynein intermediate chain attenuates HIV-1 infection and its retrograde transport^6^; 2) knockdown of dynein light chain 1^10^ or light chain 2^11^ reduce HIV-1 integrase transport or completion of reverse transcription, respectively; 3) knockdown of dynein heavy chain resulted in reduced retrograde movement of the HIV-1, viral infection, and nuclear entry^8,12^. Thus, while multiple subunits of the dynein complex are required for HIV-1 infection, cell-based experiments could not map the molecular mechanism of trafficking.

In addition to the dynein motor, previous work showed that depletion of dynactin, a dynein processivity factor, or BicD2, a cargo adaptor that connects dynein to membrane cargoes, reduces retrograde cytoplasmic trafficking of viral particles and prevents the nuclear import of the viral genome^12,13^. These findings suggest that the complex of dynein, dynactin, and the BicD2 proteins are involved in the cytoplasmic transport of HIV-1 to the nucleus but fail to show how HIV-1 inserts itself into the dynein machinery for motility.

Here, we sought to establish the molecular basis for HIV-1 retrograde motility using a purified, reconstituted system that would allow us to visualize microtubule-dependent HIV-1 trafficking *in vitro*. We discovered HIV-1 binds to the dynein motor directly instead of using a cargo adaptor such as BicD2. The direct attachment of HIV-1 to dynein allows HIV-1 to leverage microtubule-driven motility of that many divergent dynein cargo adaptors, suggesting that HIV-1 opportunistically trafficks with different cargoes in the cell. Additionally, our reconstituted system allowed us to show that 1) intact HIV-1 capsid lattice binds to the region of dynein’s tail that is comprised of dynein intermediate chain and light chains LC8 and TcTex1 and 2) teams of dynein motors drive HIV-1 motility. Finally, we create a synthetic capsid system to show that rigid cargoes, such as HIV-1 and other viral capsids, undergo reduced motility when dynein teams drive motility, which is unlike membranous cargos. Taken together, our results provide the first molecular description of viral trafficking using an *in vitro* reconstitution of HIV-1 motility. This work introduces a new paradigm of viral motility through opportunistic hijacking by direct binding to motors, allowing any cargo adaptor to support viral microtubule motility.

## Results

### HIV-1 cores bind directly to dynein for microtubule recruitment

Previous work suggested that BicD2, a dynein cargo adaptor, binds to the HIV-1 capsid or nucleocapsid *in vitro*^12,13^, which has led to the model that full-length BicD2 (BicD2^FL^) is required for the microtubule-based trafficking of HIV-1 (**Fig. 1A**). When not bound to cargo, BicD2^FL^ adopts an autoinhibited conformation involving an intramolecular interaction that prevents dynein binding^14–17^. Based on the proposed HIV-1 trafficking model, the interaction of the BicD2 cargo binding domain (coiled-coil 3) with the HIV-1 capsid releases BicD2^FL^ autoinhibition (**Fig. S1A**) to allow the N-terminus of BicD2 available to interact with dynein in a manner analogous to established BicD2^FL^ binding partners including Rab6^18^, Egalitarian mRNA^19^, and Nup358^16^.

**Fig. 1.**
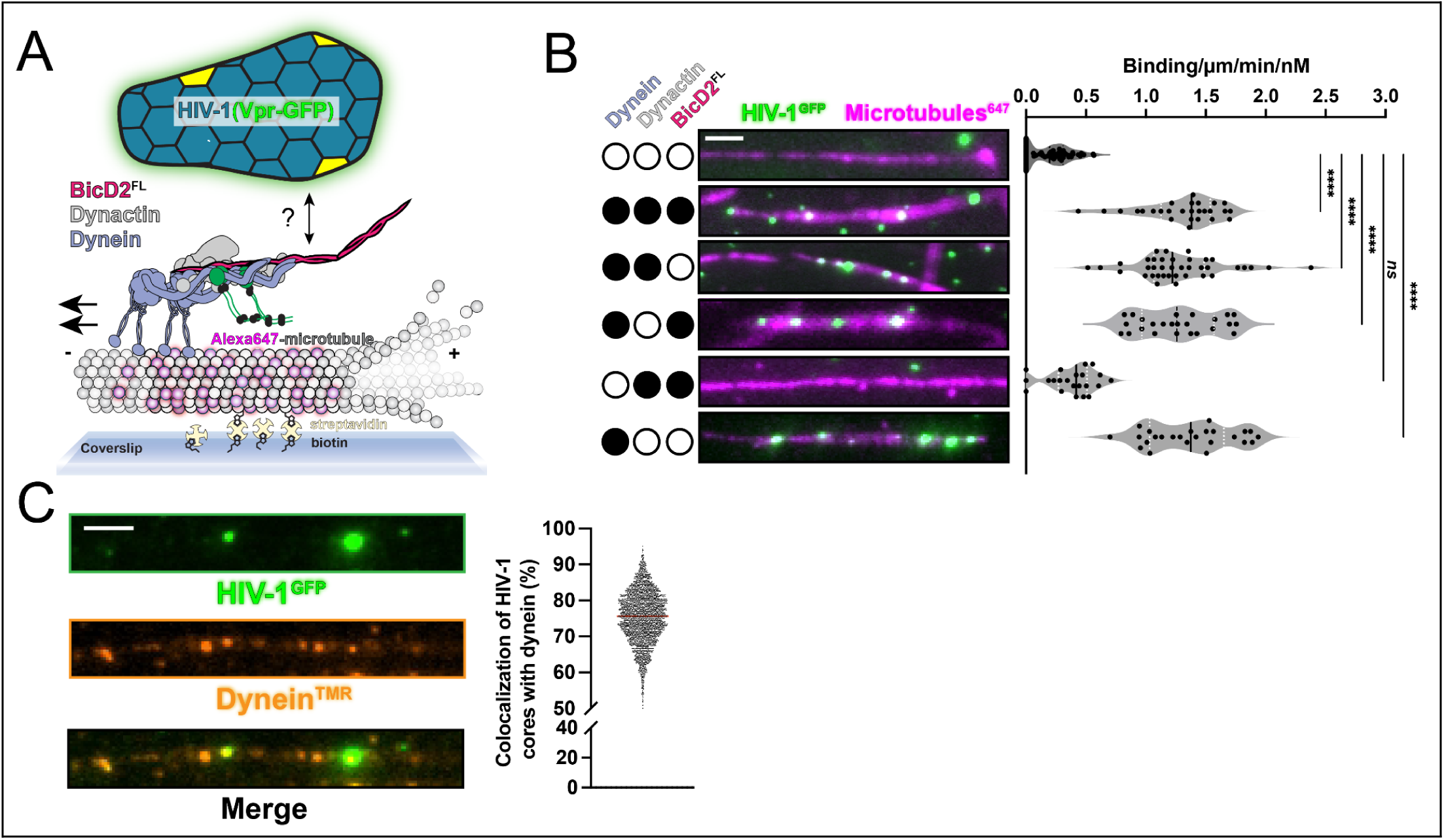
HIV-1 cores bind to dynein directly for microtubule recruitment *in vitro.* **(A)** Schematic of HIV-1 microtubule motility assay. The dynein motor assembles with dynactin and BicD2 for motility. **(B)** Microtubule recruitment assay & quantification of HIV-1 (GFP-Vpr) with dynein, dynactin, and BicD2^FL^. **(C)** Co-localization of HIV-1 (GFP-Vpr) with fluorescently-labeled dynein^TMR^. Scale bars are 5 pm. All *“* represent *P* = <0.0001 and “ns” represents *P* = 0.3991.

To directly test this model, we combined purified HIV-1 cores (fluorescently labeled with GFP-Vpr) with purified dynein, dynactin, and BicD2^FL^. HIV-1 cores are conical capsid protein (CA) surrounding an internal ribonucleoprotein complex that is competent for reverse transcription and subsequent DNA integration^20,21^. Previous work demonstrated that HIV-1 cores with GFP-Vpr incorporated allow visualization of single-core trafficking in cells^6,22,23^. After confirming the integrity of purified HIV-1 (GFP-Vpr) cores (**Fig. S1B**), we measured HIV-1 core recruitment to microtubules adhered to a coverslip via single molecule total internal reflection fluorescence microscopy (TIRF) assays (**Fig. 1A**). TIRF showed that the presence of dynein-dynactin-BicD2^FL^ increased HIV-1 cores binding to microtubules by ∼3.4 times (**Fig. 1B**). Unexpectedly, however, despite visualizing HIV-1 recruitment to microtubules, the HIV-1 cores did not undergo active motility (**Fig. S1C**).

Our observation, coupled with the fact that purified dynein is active as expected (**Fig. S2A-B**), led us to surmise that autoinhibition of BicD2^FL^ might not be released by HIV-1. If this assumption was true, it suggested that the BicD2^FL^ cargo binding domain does not interact with the HIV-1 core. To test this hypothesis, we designed drop-out experiments using single-molecule TIRF to determine the minimal components needed for HIV-1 microtubule recruitment. Interestingly, removing BicD2^FL^ and/or dynactin from the imaging reaction did not affect HIV-1 core recruitment to microtubules (**Fig. 1B** and **Fig. S1C**). In contrast, removing dynein showed a dramatic loss of HIV-1 core microtubule recruitment (**Fig. 1B** and **Fig. S1C**). These data indicate that dynein binds directly to HIV-1 cores without needing the cargo adaptor BicD2^FL^. To directly visualize the interaction, we imaged HIV-1 cores with fluorescently labeled dynein and saw that 75.6% of microtubule bound HIV-1 cores are colocalized with fluorescently labeled dynein (**Fig. 1C**). Thus, HIV-1 binds to dynein directly for microtubule recruitment *in vitro*.

### HIV-1 exploits multiple dynein-activating adaptors for motility

Our results suggest that dynein-dynactin-BicD2^FL^ does not support the motility of HIV-1 because HIV-1 binds to dynein directly instead of binding and activating BicD2^FL^. This model predicts that dynein activation is independent of HIV-1 binding. To test this, we compared HIV-1 motility with dynein and dynactin using either BicD2^FL^ or an N-terminal truncation of BicD2 that is missing the autoinhibitory cargo binding domain and constitutively activates dynein motility (**Fig. S2A**)^24,25^. We found that dynein-dynactin incubated with BicD2^1–598^ drove the processive motility of HIV-1 cores (**Fig. 2A**). Moreover, these results indicate that the BicD2 C-terminal cargo binding domain is not needed for HIV-1 binding to microtubules or motility, as previously proposed^12,13^.

**Fig. 2.**
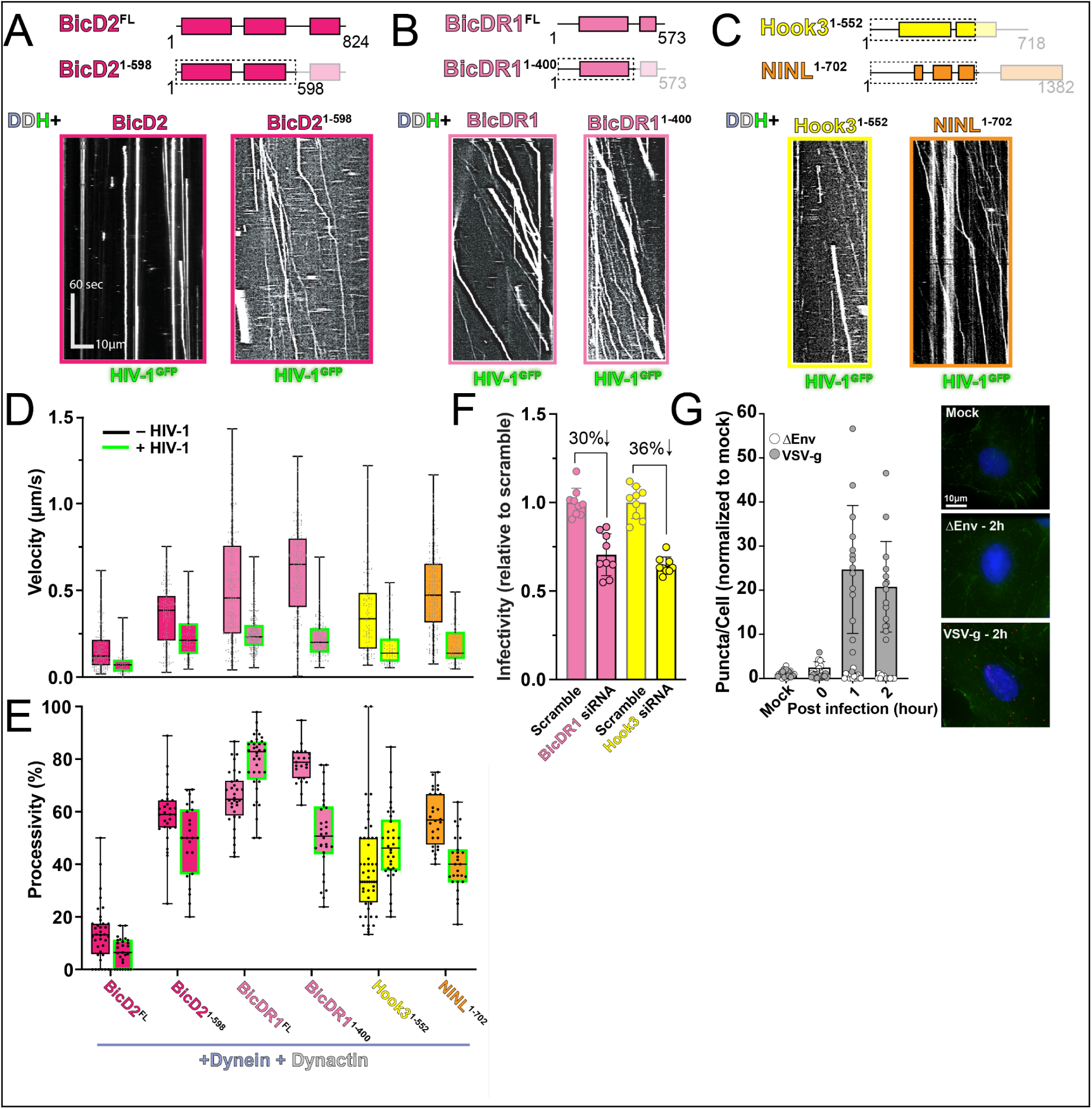
HIV-1 exploits multiple dynein-activating adaptors for *in vitro* motility. Schematic coiled-coil presentation of BicD2 **(A),** BicDRI **(B),** Hook3, and NINL **(C)** constructs shown alongside reconstitution of HIV-1 motility with adaptor and dynein-dynactin. **(D)** Quantification of velocity for dynein-dynactin-adaptor and HIV-1 cores. (E) Quantification of processivity for dynein-dynactin-adaptor and HIV-1 cores. (F) HIV-1 infection assays using luciferase reporter for siRNA-mediated knockdowns of BicDRI and Hook3 in A549 cell. (G) Proximity ligation assay between HIV-1 CA (P24) and BicDRI for uninfected (Mock), negative control HIV-1 strain that cannot enter cells (AEnv), and HIV-1 pseudotyped with VSV-g Env protein. (Left) Quantification of puncta/cell. (Right) Representative fluorescence images for 2 hours post infection.

Considering that BicD2^1–598^ activates HIV-1 motility without a cargo binding domain, we hypothesized that any cargo adaptor promoting dynein motility would enable HIV-1 motility. To test this, we incubated dynein and dynactin with several cargo adaptor constructs previously shown to activate dynein motility *in vitro* and asked if the resulting complexes could generate processive HIV-1 motility. Here, we used full-length and truncated versions of a BicD2 ortholog which is not autoinhibited, called BicDR1 (BicDR1^FL^ and BicDR1^1–400^)^26,27^, and two well-characterized truncations of Hook3 (Hook3^1–552^)^28,29^ and Ninein-like (NINL^1–702^)^30^ **(Fig. S2A-B)**. Importantly, none of these cargo adaptors have previously been implicated in HIV-1 trafficking or infection. Surprisingly, we observed processive motility of HIV-1 with all non-autoinhibited cargo adaptor constructs tested (**Fig. 2A-C**). This finding supports our model that dynein-mediated HIV-1 motility does not require HIV-1 binding to adaptors as cargo. Additionally, we observed almost the same landing rate of HIV-1 on microtubules with all adaptors, which suggests that the identity of the cargo adaptors does not alter the basal association of HIV-1 and dynein (**Fig. S2C**).

Remarkably, we found that dynein-driven HIV-1 motility is significantly slower than the motility of dynein alone (**Fig. 2D**). Analysis of the velocity of HIV-1 bound to all cargo adaptors showed an average velocity of ∼0.16 µm/s, which is 2-4X slower than the overall velocity for dynein-dynactin with the same cargo adaptors (without HIV-1 cargo) (**Fig. 2D**). Velocity of dynein-mediated HIV-1 movement that we measured *in vitro,* however, is in agreement with the velocity measured in cells (∼0.1-0.3 μm/sec)^31^.

While we did not observe a significant difference in the velocity of HIV-1 being trafficked by dynein-dynactin complexed to different cargo adaptors, we did observe a difference in processivity with various adaptors (**Fig. 2E**). Over 80% of the HIV-1 cores that were recruited to the microtubule move processively in the presence of dynein-dynactin-BicDR1^FL^, whereas this was reduced to ∼50% and less for the N-terminal truncations of BicDR1, BicD2, Hook3, and NINL. Finally, <10% of the HIV-1 recruited to the microtubule via dynein bound to BicD2^FL^ was processive. The observed processivity trends are directly correlated to the intrinsic ability of each cargo adaptor to activate dynein-dynactin in the absence of HIV-1, showing that HIV-1 does not affect the processivity of dynein activated by divergent cargo adaptors (**Fig. 2E**).

Our model suggests that any cargo adaptor that enables dynein motility can function as a host factor that supports HIV-1 infection. To test this model, we depleted BicDR1 and Hook3 cargo adaptors from A549 cells and monitored infectivity. We did not test NINL in the infectivity assays because NINL contributes to innate immunity via interferon signaling^32^. In support of our new model of HIV-1 motility, the knockdown of BicDR1 or Hook3 in A549 cell lines reduced HIV-1 infectivity by nearly 30% and 36%, respectively (**Fig. 2F, Fig. S3A-C**). And infectivity was reduced by 42% for BicDR1 depleted CHME-5 cell lines (**Fig. S3D-E**). Interestingly, we observed within Hela TZM-bl cells no change in HIV-1 infectivity following knockdown of either BicDR1 or Hook3 (**Fig S3F-G**). These results recapitulate previous studies and suggest a cell-type-specific utilization of dynein-activating adaptors^12,13^.

Host factors that support infection should also colocalize with HIV-1. Using proximity ligation assay to monitor the proximity of HIV-1 capsid with BicDR1 in A549 cells, we found an ∼80x increase in HIV-1-BicDR1 colocalization 1-2 hours post-infection (**Fig. 2G and Fig. S3H)**. This is comparable to the colocalization previously observed with BicD2^13^. Together, our work suggests that HIV-1 exploits multiple dynein cargo adaptors to support infection.

### The HIV-1 capsid is sufficient for dynein-directed motility

After observing that HIV-1 cores can bind to dynein and undergo directional motility, we hypothesized that dynein is binding to the viral capsid shell of the cores. The capsid shell surrounding cores binds to numerous host factors^33^ and remains nearly intact when docked at the nuclear pore^34–36^, thus providing a potential binding platform for interaction with dynein. To test whether the capsid protein (CA) can bind dynein directly, we performed TIRF motility assays using cross-linked, fluorescently-labeled capsid particles produced by self-assembly of recombinant CA (A92E/A204C) under conditions that yield a mixture of capped tubes and cones from building blocks of CA hexamers and pentamers (**Fig. 3A, Fig. S4A**). We saw that, like HIV-1 cores, dynein alone recruits CA particles to microtubules, and BicD2^FL^ is unable to activate motility (**Fig. 3B**). Importantly, when constitutively active cargo adaptors were included in the assay, we observed robust motility for all adaptors tested (BicD2^1–598^, BICDR1^FL^, BicDR1^1–400^, HOOK3^1–552^, and NINL^1–702^) (**Fig. 3B**). Again, similar to HIV-1 cores, we see that the velocity of moving CA particles is reduced compared to dynein not loaded with viral cargo (**Fig. 3C**). Additionally, the overall dynein-dependent microtubule binding rate of CA-particles is similar to that observed with HIV-1 cores (**Fig. 3D**). Based on these results, the intact HIV-1 capsid lattice is sufficient for binding motile dynein *in vitro*.

**Fig. 3.**
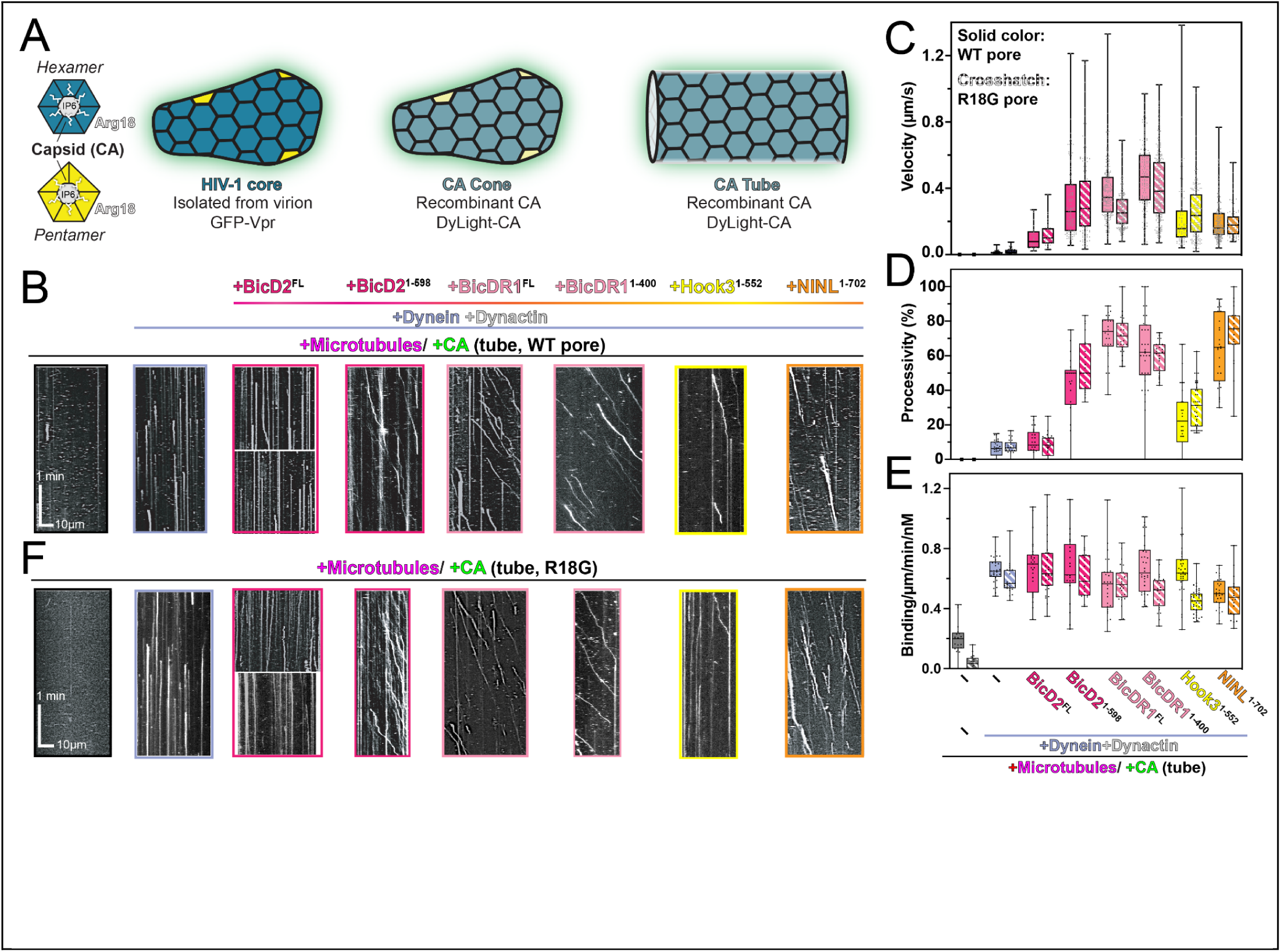
HIV-1 capsid is sufficient for dynein-directed motility *in vitro.* **(A)** Structural models of capsid (CA) hexamers and pentamers shown alongside HIV-1 cores, CA tubes (formed with high salt or A92E/A204C), or CA cones (formed by IP6 or A92E/A204C). (B) Microtubule recruitment and motility reconstitution of CA particles (A92E/A204C) with dynein, dynactin, and different activators. Quantification of velocity **(C)** processivity **(D)** and recruitment (E) for HIV-1 tubes. **(F)** Motility of CA tubes with mutated pores (R18G) with quantification in (C), (D) & (E).

Two well-established and discrete sites on the HIV-1 capsid lattice are the central pore of the hexamer and the FG-binding pocket (**Fig. S4B**). Given that each site interacts with multiple, unique host factors, we speculated that dynein may engage with the viral capsid via one of these sites.

First, we examined if dynein interacts with the central pore, which lies at the center of CA hexamers and is lined by six positively charged arginine residues (R18) (**Fig. 3A, Fig. S4B**). The R18 pore binds to nucleotides^37^ and IP6^38,39^, where the latter plays a critical role in HIV-1 capsid maturation^38^. Recent work also showed that the kinesin-1 adaptor, FEZ1, binds the central pore of hexamers^40^. To test if dynein binds to the R18 pore, we purified a mutant CA particle with glycine introduced into the pore (R18G). In TIRF assays, capsid particles assembled from R18G CA (**Fig. S4A**) were recruited to microtubules and transported similarly to capsid particles with R18 (**Fig. 3E**), with nearly the same velocity (**Fig. 3C**) and binding rate as wildtype (**Fig. 3D**). The wild-type pore capsid, however, showed a slightly higher binding rate to the microtubules even in the absence of dynein (**Fig. 3D**). We reason that this is due to a partial electrostatic interaction of the positively charged pores in wildtype with the negatively charged surface of microtubules. Additionally, IP6 (which should act as a competitive inhibitor to any factor that binds to the R18 pore) did not affect the microtubule association, processivity, or velocity of capsid particles (**Fig. S4D**). From these data, we conclude that HIV-1 does not bind dynein using the central pore.

Next, we examined if dynein binds to the FG-binding pocket by monitoring dynein, dynactin, BicDR1 (DDR)-mediated trafficking of HIV-1 cone-shaped capsid (**Fig. S4C)** in presence of PF74. Small molecules such as GS-6207 and PF74^41^ and host factors such as Nup153 and CPSF6 bind at the FG-binding pocket (**Fig. S4B)**^42^. PF74 in excess of HIV-1 CA particles did not affect HIV-1 capsid binding to microtubules, processivity, or velocity (**Fig. S4D**). Altogether, these results suggest that dynein does not bind to either the central pore or the FG-binding pocket of HIV-1 capsid.

### HIV-1 capsid binds to the tail domain of dynein

Human cytoplasmic dynein is a ∼1.4 MDa dimeric complex consisting of a heavy chain (DHC, ∼530 kDa) and several associated subunits: the intermediate chain (DIC), the light intermediate chain (DLIC); and three light chains (DLC), Rob1/LC7, LC8, and Tctex1 (**Fig. 4A**). The C-terminal two-thirds of the heavy chain form the ATP-driven motor domain (∼380 kDa) while N-terminal part of the heavy chain associates with DIC and DLIC, where the DLCs bind to the DIC subunit^3,43^. To determine which regions or subunits of dynein contribute to HIV-1 binding, we generated multiple constructs of dynein that have specific regions truncated or accessory chains deleted and monitored the ability of HIV-1 to engage with each construct.

**Fig. 4.**
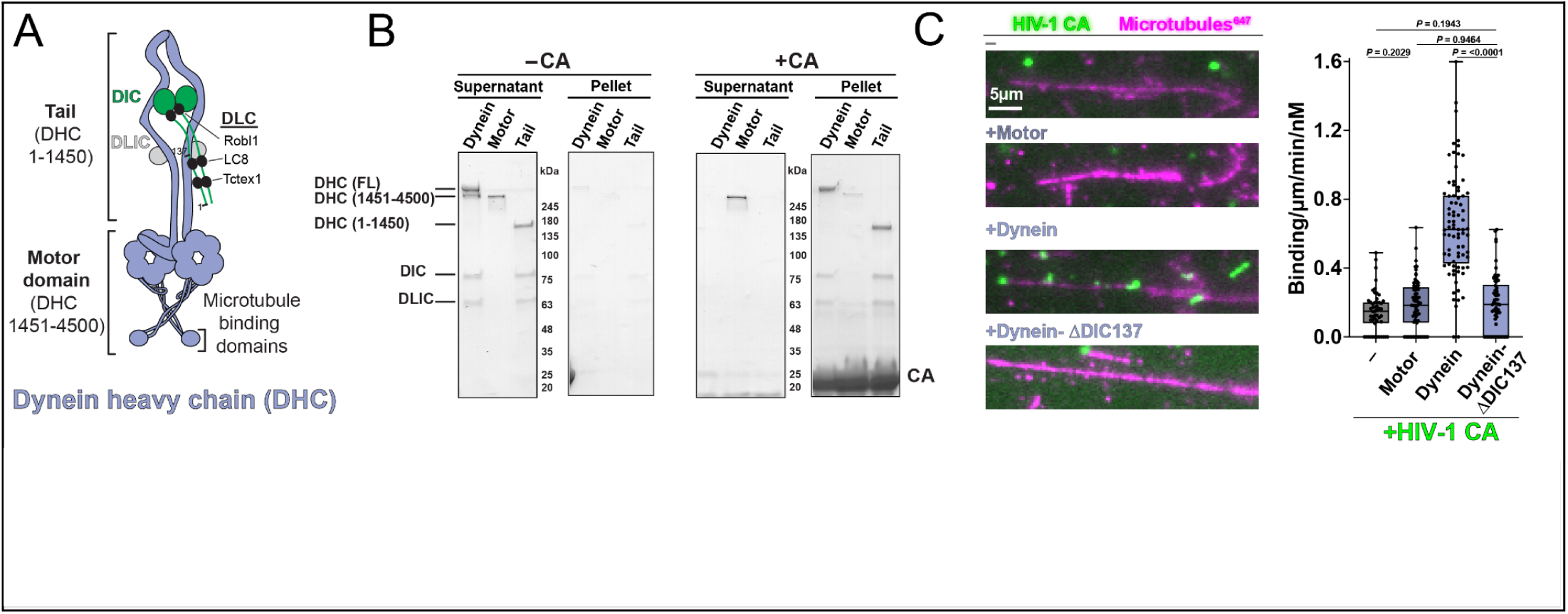
The HIV-1 capsid binds to the dynein tail domain. **(A)** Schematic of dynein structure and segments. **(B)** Co-pelleting of CA tubes (14C/45C) with dynein, dynein motor and dynein tail. **(C)** HIV-1 microtubule recruitment assay using motor domain, full-length dynein, and dynein-ADIC137 with CA tubes (A92E/A204C capsid).

First, we tested whether cross-linked HIV-1 CA tubes (A14C/E45C) bind dynein’s tail or motor domain. To do this, we purified two distinct constructs of dynein: one that consisted of only the tail region (DHC N-terminus, DIC, DLIC, and DLCs)^26^ and another construct that contained only the ATPase motor domain (DHC 1451-4500) **(Fig S5A-B)**. Using these two constructs, we performed a co-sedimentation experiment with CA tubes *in vitro*. When co-pelleted with cross-linked CA tubes (A14C/E45C), we saw robust binding for full-length dynein and the tail, whereas the motor domain showed little binding (**Fig. 4B**). Thus, capsid binds dynein’s tail and not the ATPase motor domains.

To confirm the cosedimentation results and extend our binding data, we utilized HIV-1 microtubule recruitment assays with dynein truncation constructions. Here, we compared HIV-1 CA tube microtubule recruitment using full-length dynein, motor domain only (DHC 1451-4500), and Dynein-ΔDIC137. Previous work has implicated dynein light chain 1 (LC8) as a factor that is required for HIV-1 replication^11^ and, in some other retroviruses, LC8 interacts with the capsids^44,45^. LC8 is missing from the Dynein-ΔDIC137 complex, as is TcTex1 (**Fig. S5C-D**). First, we showed they bound microtubules in a comparable manner in the absence of CA-tube or capsid (**Fig. S5E**). Next, we monitored dynein-mediated microtubule recruitment of fluorescent CA particles (A92E/A204C) with each dynein construct. Consistent with the sedimentation data generated with CA tubes (A14C/E45C), we observed that the motor domain only (DHC 1451-4500) did not recruit HIV-1 CA particles to microtubules. In contrast, full-length dynein recruited CA particles to microtubules, confirming the co-sedimentation data (**Fig. 4C**). Interestingly, we also observed a dramatic loss in CA particle recruitment to microtubules when we used Dynein-ΔDIC137 (**Fig. 4C**).

This data suggests that the HIV-1 CA lattice binds to dynein’s tail domain utilizing the N-terminus of dynein intermediate chain with LC8 and TcTex1.

### HIV-1 recruits a team of dynein motors for motility

Throughout our studies, we observed significantly reduced velocities of motile HIV-1 compared to the velocity of the dynein-dynactin-cargo adaptor alone (**Fig. 2D**). To examine the parameters affecting the velocity of HIV-1 bound dynein, we monitored HIV-1 motility in the presence of TMR-labeled dynein and compared fluorescence intensities for individually walking dynein-dynactin-BicDR1^FL^ motors versus dynein colocalized with HIV-1 cores (**Fig. 5A**). Our analysis showed that the fluorescence was on average 7.2X brighter for dyneins associated with HIV-1 motility (**Fig. 5B**). Thus, moving HIV-1 recruits multiple dynein motors.

**Fig. 5.**
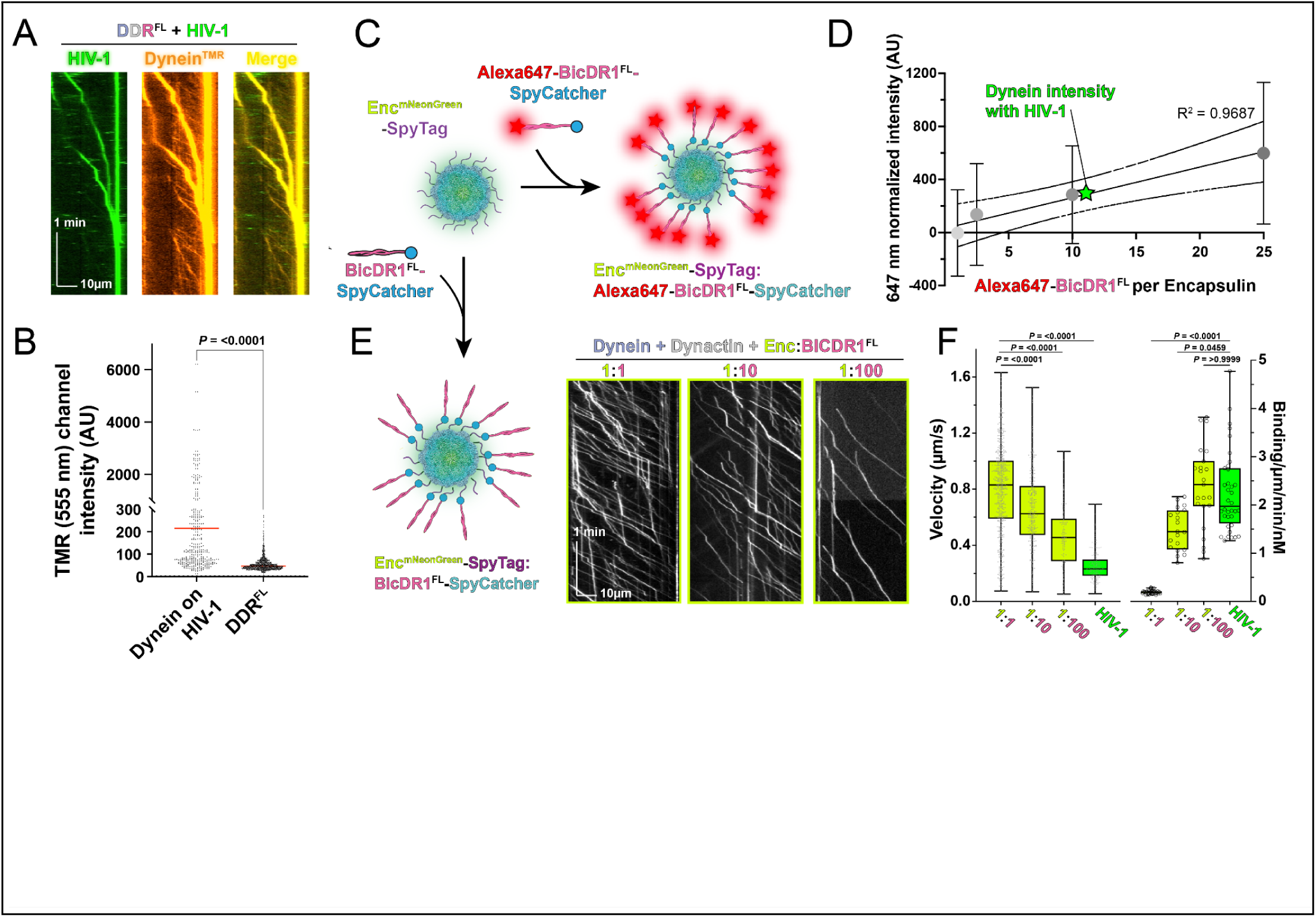
HIV-1 recruits a team of dynein motors for motility. **(A)** Representative kymographs of the colocalization of HIV-1 cores (Vpr-GFP) and TMR-labeled dynein in moving DDR^FL^-HIV-1 complexes. Each channel is shown separately (left and middle) and merged (right). **(B)** Related to panel A, florescence intensity of TMR-555 labeled dynein in single DDR vs. co-localized with HIV-1. A random frame out of 4 min movie was selected for quantification by Imaged ComDet plugin. **(C)** Schematic design of Encapsulin-SpyTag fluorescently labeled with SNAP-BicDR1^FL^-SpyCatcher at increasing ratios of BicDR1^FL^:Encapsulin. (D) Comparison of BicDR1^FL^:Encapsulin fluorescence standard curve versus fluorescent signal of dynein on HIV-1 cores. (E) Formation of BicDR1^FL^-SpyCatcher-labeled Encapsulin complexes and representative kymographs of Encapsulin -mNeonGreen signal with different ratios of BicDRI in the presence of dynein and dynactin. **(F)** Quantification of velocities and binding rates for encapsulin complexes at each BicDRI ratio shown alongside data of HIV-1 core with dynein, dynactin and BicDRI.

To quantify how many dynein motors are bound to HIV-1 cores, we compared dynein fluorescence to a standard curve generated using engineered encapsulins. Encapsulins are customizable bacterial nano-compartments that can be exploited for biophysical assays, such as genetically encoded multimeric nanoparticles (GEMs)^46^. The encapsulin construct used in this study assembles into a T=4 (240 protomers, 42 nm diameter) icosahedral protein cage^47^, encapsulating mNeonGreen via an N-terminal fusion to its natively loaded cargo protein (IMEF)^48,49^. Each encapsulin subunit has a surface-exposed C-terminal SpyTag that can efficiently form a covalent bond with its 10 kDa partner protein SpyCatcher (**Fig. S6A**)^50^.

Fluorescently-labeled SNAP^Alexa647^–BicDR1^FL^-SpyCatcher was mixed with Encapsulin-SpyTag at specific ratios to create decorated encapsulins with 1, 2.5, 10, 25, or 100 SNAP^Alexa647^–BicDR1^FL^-SpyCatcher proteins per encapsulin (**Fig. 5C, S6B-C**). The fluorescence intensity of each SNAP^Alexa647^–BicDR1^FL^-SpyCatcher:Encapsulin-SpyTag was measured by TIRF on an untreated coverslip (**Fig. S6D**). A standard curve was generated by plotting the mean fluorescence emission of each construct against the number of dyes on each encapsulin (SNAP^Alex647^–BicDR1^FL^-SpyCatcher) (**Fig. S6F-G**). In parallel, SNAP-dynein was labeled with the same Alexa647 dye and allowed to bind to HIV-1 cores at saturating concentrations. SNAP^Alexa647^–Dynein-bound HIV-1 complexes were adhered to untreated coverslips and imaged using 647 nm excitation (**Fig. S6E**) and the average dynein intensity was calculated from dyneins colocalized with HIV-1 cores (**Fig. 5E**). After comparing dynein intensity to the encapsulin standard curve, we see that 11.58 ± 11.27 dynein dimers bind per HIV-1 core. The broad distribution of this data suggests a significant variation of dynein motors per HIV-1.

We hypothesized that the reduced velocity we observed was due to the negative interference among multiple dyneins bound to the surface of the HIV-1 capsid. It is possible that collective yet uncoordinated stepping would result in a reduction in velocity^51^. This is surprising given that membranous cargo, like phagosomes and endosomes, recruit 10-15 dynein motors in cells and still move with a relatively fast velocity (2μm/sec)^52^. We speculated that the rigidness of the cargo could cause a difference in dynein behavior on membranes versus HIV-1. In fact, there has been no systematic study into how multi-dynein motors function on rigid cargoes the size of HIV-1.

To determine if rigid cargo prevents the coordinated stepping of multiple dynein motors, we utilized the rigid, synthetic Encapsulin-SpyTag cargo. To do this, we bound increasing numbers of BicDR1^FL^-SpyCatcher to assembled Encapsulin-SpyTag, and imaged the complexes using the incorporated mNeonGreen signal within the Encapsulin in the presence of saturating concentrations of dynein and dynactin (**Fig. 5E**). We found that increasing the ratio of BicDR1^FL^-SpyCatcher per encapsulin (1:1→ 1:10→ 1:100), decreases the overall velocity while increasing the encapsulin complexes’ landing rate (**Fig. 5F**). This result indicates that rigid cargoes such as encapsulin or HIV-1 capsid with multiple bound dyneins have reduced velocities compared to membranous cargo due to inter-motor interference during trafficking.

## Discussion

Here, we utilized in vitro reconstitution to discover that HIV-1 hijacks the dynein transport machinery for microtubule motility. Unexpectedly, we see that dynein binds via its tail domain to the HIV-1 capsid and separately recruits an adaptor protein for activation, allowing HIV-1 to utilize multiple dynein cargo adaptors for motility. Finally, we show that HIV-1 displays reduced velocity compared to native cargoes because teams of dynein motors recruited to the capsid display negative interference with respect to velocity. This work establishes a new model of HIV-1 motility and shows that HIV-1 can bind to motor proteins directly to hijack host cell transport machinery.

Our data supports a new model of HIV-1 trafficking where HIV-1 is an opportunistic hijacker of dynein motility (**Fig. 6**). In this model, HIV-1 directly binds to the dynein motor instead of using a cargo adaptor. In our *in vitro* system, we observed that HIV-1 could bind multiple dynein motors without the need of additional cargo adaptors or dynactin (**Fig. 6A**). Interestingly, when HIV-1 was bound to dynein, there was no motility of the HIV-1:dynein complex until constitutively active cargo adaptors were added. This may represent a way for HIV-1 to localize to microtubules.

**Fig. 6.**
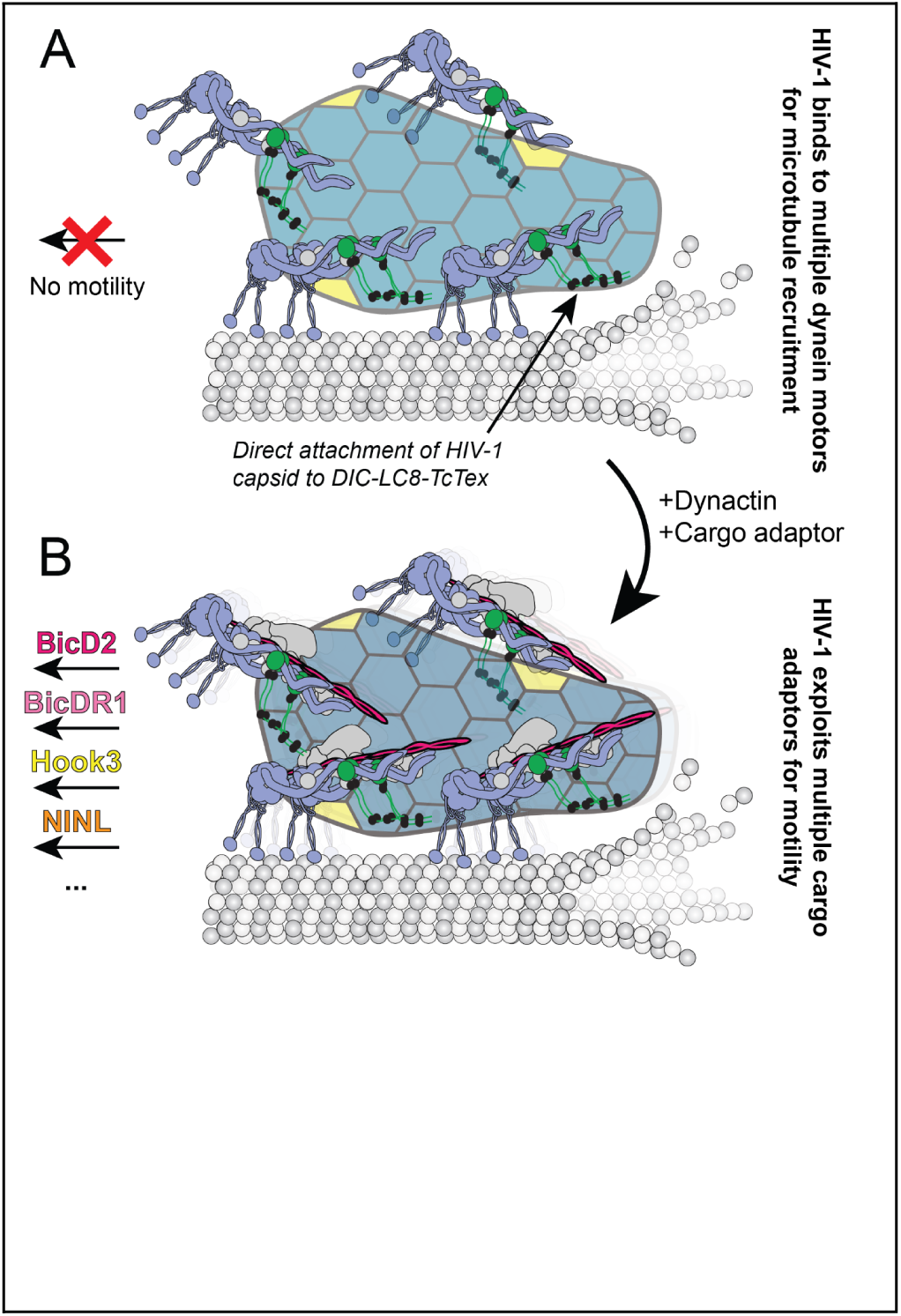
Model of opportunistic hijacking of dynein motor proteins by HIV-1. **(A)** HIV-1 recruitment to microtubule via multiple dynein motor proteins. The capsid of HIV-1 binds to the tail domain of dynein using the composite binding site of DIC-LC8-TcTex1. When bound to dynein alone, the HIV-1 :dynein complexes are immotile. (B) HIV-1 achieves microtubule motility by exploiting active dynein cargo adaptors in conjunction with the dynactin core regulator.

When dynactin and cargo adaptors are present with HIV-1 and dynein, we saw motility of both HIV-1 cores and HIV-1 capsid assemblies (**Fig. 6B**). Importantly, given that HIV-1 is bound to dynein directly, any dynein cargo adaptor could support HIV-1 motility. This result suggests that HIV-1 is agnostic to the cargo adaptor for motility and will opportunistically hijack motility of cargoes in cells. Further work is needed to place these data into the context of cargos such as Rab6-vesicles which have previously been shown to activate dynein motility via BicD2^18^.

Unexpectedly, we observed reduced velocity for HIV-1 complexes on microtubules compared to activated dynein motility. Previous work with active dynein alone^24,25^, with vesicles *in vitro*^18^, or purified dynein-driven organelles^52^ show that dynein motility is ∼1-2 μm/sec. In contrast, HIV-1 motility for the adaptors tested showed velocities between 0.1 - 0.4 μm/sec, suggesting that dynein driven motility for HIV-1 is different than previously studied complexes. To explore this, we exploited a novel engineered encapsulin system to show that rigid cargoes such as HIV-1 exhibit negative inference between motors when the numbers of active dyneins are increased per rigid cargo. These results suggest that rigid cargoes, such as viruses, experience different motility than vesicles which may play a role in viral trafficking and detection in the cytoplasm by the innate immunity detection machinery.

Our reconstitution of HIV-1 trafficking via dynein represents the first example of viral trafficking using reconstituted components, allowing us to dissect the nature of the interaction between HIV-1 and dynein. Previous studies implicated direct binding of viral subunits to dynein motor proteins^53,54^, such as the Adenovirus hexon submit binding to dynein^55^. Our system allowed us to visualize HIV-1 direct binding to dynein and exploiting multiple cargo adaptors for motility, demonstrating a new mode of viral trafficking by utilizing any dynein cargo adaptor for movement.

More broadly, our work shows that viruses such as HIV-1 can bypass interaction with a cargo adaptor by directly binding to motor proteins. Thus, direct motor protein hijacking may represent an important mode of trafficking for other viruses that utilize dynein and kinesin motor proteins.

## Acknowledgments

We thank all members of the Cianfrocco Laboratory for their support and discussions, especially Dr. Anthony Ludlam for dynactin purifications. We also thank the Microtubule and Motor Protein community at the University of Michigan, which comprises the Cianfrocco, DeSantis, R. Ohi, Sept, and Verhey labs. We thank Dr. Andrew Carter for the Dynein-ΔDIC137 baculovirus plasmid. We are grateful for discussions with the Pittsburgh Center for HIV-1 Protein Interactions, Christian Freniere and Dr. Yong Xiong, Drs. Barbie Ganser-Pornillos and Owen Pornillos, and Drs. Allison Gicking and Will Hancock. We recognize the following grants that supported this work: P50AI150481 (SB & MAC), R21AI152869 (SB & MAC), R01AI162694 (DL & EMC), R35GM133325 (MPA & TWG), R35GM146739 (JPG & MED), R21AI152869 (JS & CA), NSF 2142670 (JPG & MED), and APP1182212 (WP & TB). The University of Michigan Cryo-EM Facility (U-M Cryo-EM). U-M Cryo-EM is grateful for support from the U-M Life Sciences Institute and the U-M Biosciences Initiative.

## Author contributions

Conceptualization: SB, MAC

Investigation: SB, DL, MPA, JPG, WP, JS

Funding acquisition: MAC, EMC, TWG, TB, CRA, MED

Supervision: MAC, EMC, TWG, TB, CRA, MED

Writing - original draft: SB, MAC

Writing - review & editing: SB, DL, MPA, JPG, WP, JS, MED, CRA, TB, TWG, EMC, MAC

## Declaration of interests

Authors declare that they have no competing interests.

## Data and materials availability

All plasmids and data are available upon request.

## Methods

### Protein expression, purification, and labeling

#### Dynein and Dynactin purification

Bovine dynactin from cow brain and recombinant human cytoplasmic dynein expressed in insect cells were purified as described before ^56^.

Human cytoplasmic dynein tail (DHC 1-1450 with accessory proteins DYNC1I2, DYNC1LI2, DYNLT1, DYNLL1, and DYNLRB1), human cytoplasmic dynein motor domain (DHC 1451-4500) and dynein-ΔDIC1-137 were all recombinantly expressed in insect cells. In summary, the baculovirus was prepared for dynein tail segments (pACEBac1 donor vector), dynein motor domain (pFastBac donor vector) and dynein-ΔDIC137 (DHC full length, with accessory proteins DYNC1I2 (138-640), DYNC1LI2, DYNLT1, DYNLL1, and DYNLRB1) in SF9 cells using standard methods. The proteins were expressed in SF9 or High Five™ Cells (Thermo Fisher) (BTI-TN-5B1-4) using passage one of viruses. Lysis buffer includes 50 mM HEPES pH 7.4, 100 mM NaCl, 10% glycerol, 0.5 mM EGTA, 1 mM DTT, 0.1 mM ATP-Mg, 1 unit Benzonase, SIGMAFAST™ Protease Inhibitor Tablets + 0.5 mM Pefabloc. The proteins were bound to IgG Sepharose 6 Fast-Flow (GE Healthcare) via the ZZ tag on the heavy chain. After washing with lysis buffer and DynBac TEV buffer (50 mM Tris-HCl pH 8, 2mM Mg-Acetate, 1mM EGTA, 250mM K-Acetate, 10% glycerol, 1 mM DTT, 0.1 mM ATP-Mg), proteins were eluted by treating the beads with at home-purified TEV protease in DynBac TEV buffer. The eluent was subjected to size-exclusion chromatography using GF150 buffer (25 mM HEPES pH 7.4, 150 mM KCl, 0.5 mM EGTA, 1 mM DTT). The collected fractions were concentrated and stored in a GF150 buffer including 10% glycerol at −80 °C. SDS-PAGE followed by coomassie or silver staining was used to evaluate the purity, size and the presence of the related polypeptide chains for each construct.

To analyze microtubule recruitment, the SNAP tag at the N-terminus of the heavy chain in each construct was used for fluorescence labeling by SNAP-Cell® TMR-Star or SNAP-Surface Alexa Fluor® 647. The extra dye was removed by size-exclusion chromatography. The efficiency of labeling was determined by nanodrop and based on the protein concentration.

#### Full-length and truncated BicD2 and BicDR1

8xHis-zz-TEV-mBICD2 (full-length or truncated 1-560) was codon optimized for expression in SF9 insect cells and synthesized by Genscript. The genes were subcloned into pFastBac using NEBuilder HiFi DNA assembly (NEB E2621S). zz-TEV-Halo-hBicD2^FL^-6xHis in donor vector pKL for expression in insect cells and zz-TEV-Halo-hBicD2(1-598) in pET28a vector for expression in *E. coli* were received from DeSantis Lab.

Baculoviruses were prepared from pFastBac or pKL (for Halo-BicD2^FL^) donor vectors using standard methods. SF9 cells at 2 × 10^6^/ml density were infected with passage 2 of virus at 1:100 ratio and harvested 48 h later. Each gram of cell pellet was resuspended in 8 ml of buffer A (HEPES-KOH 30 mM pH 7.4, KCl 150 mM, Glycerol 10%, EGTA 1 mM, 10 mM BME) in the presence of PMSF 0.1 mM, SIGMAFAST™ protease inhibitor and 25 units benzonase. The cells were lysed, and the lysate was cleared as described for dynein. The clear lysate was incubated with washed IgG Sepharose Fast Flow beads (Cytiva) at 4 °C for >2 hours to be purified via their ZZ tag. The BicD2 bound beads were washed with 100 ml buffer A, including 0.02% NP40, followed by 100 ml wash with TEV buffer (Tris 10mM, pH 8, KCl 150 mM, 1 mM EGTA, 1mM DTT, 10% Glycerol). The beads were resuspended in 0.5-1 ml of TEV buffer, and incubated with 100 µg/ml of in-home purified TEV protease overnight. The eluent was subjected to further purification with size-exclusion chromatography using Superose 6 300/10 in the presence of GF150 and finally stored in a low-salt buffer (25 mM HEPES pH 7.4, 50 mM KCl, 0.5 mM EGTA, 5% glycerol).

The Halo-hBicD2 N-terminal domain (1-598) in the pET28a vector was expressed in *E*. *coli* using HyperBroth media induced with 0.5 mM IPTG. Cells were suspended and sonicated in lysis buffer (HEPES 30mM pH 7.4, NaCl 100 mM, Mg-acetate 2mM, EGTA 0.5mM, glycerol 10%) in the presence of 1mg/ml of lysozyme, 1 mM DTT, and SIGMAFAST™ protease inhibitor. The lysate was cleared by a 45-minute centrifugation at 100,000 x g at 4 °C. IgG beads washed and equilibrated with the lysis buffer were added to the lysate. After a 2-hour incubation, the beads were collected and washed with lysis buffer + DTT 1mM, followed by 100 ml wash with TEV buffer (50 mM Tris-HCl pH 8, 2mM Mg-Acetate, 1mM EGTA, 250mM K-Acetate, 10% glycerol, 1 mM DTT). The beads were suspended in 0.5-1 ml TEV buffer and incubated with TEV protease at 100 µg/ml at 4 °C overnight. The eluted protein was further purified by size-exclusion chromatography using GF150. The fractions of Halo-hBicD2 (1-598) were combined, concentrated, and re-buffered into GF150, 1mM DTT and 10% glycerol. Halo-BicD2 (FL or truncated) was labeled by HaloTag® Alexa Fluor® 660 Ligand (Promega) by mixing protein solution with the dye at 1:2 molar ratio for 30-60 min at 4 °C. Excess dye was removed by size-exclusion chromatography.

Hisx8-ZZ-TEV-SNAP-mBicDR1^FL^ in pOmnibac vector was received from Addgene (catalog #: 111859 and 111858). N-terminal truncated BicDR1 (1-400) was prepared from the full length BicDR1 in the same vector by NEBuilder HiFi DNA Assembly. Baculoviruses were prepared using SF9 insect cells as described before. Passage 1 of viruses was used to express the proteins in High Five™ Cells (Thermo Fisher). The cells harvested 48 hours post inoculation were lysed in 50 mM Tris.HCl pH 8, 150 mM NaCl, 10% Glycerol, 1 mM EGTA, 4 mM DTT supplemented with 0.2% NP-40 and Benzonase in presence of protease inhibitor (SIGMAFAST™ and 0.1 mM PMSF). Clear lysates were incubated with IgG Sepharose Fast Flow beads (Cytiva) as described for BicD2 purification. Proteins were eluted from IgG beads using TEV protease (at 100 µg/ml) in TEV buffer (Tris.HCl pH 8 25 mM, NaCl 50 mM, 10% glycerol) after incubation at 16 °C for 1-2 hours. For fluorescence labeling, eluated proteins from IgG beads were centrifuged at 17,000 x g for 45 minutes, and 500 µl of each protein were incubated with SNAP-Cell® TMR-Star 555 in 1:3 molar ratio at 4 °C for 2 hours. The labeled and unlabeled protein samples were further purified by size exclusion chromatography using Superose 6 300/10 equilibrated with Tris.HCl 50 mM pH 8.0, NaCl 50mM, and EGTA 1mM.

#### SpyCatcher BicDR1

The ZZ-BicDR1^FL^-SpyCatcher and ZZ-SNAP-BicDR1^FL^-SpyCatcher constructs in pFastBac donor vector were made by gibson assembly using NEBuilder HiFi DNA Assembly. The Baculoviruses were prepared using SF9 cells. High Five™ Cells (Thermo Fisher) at 2 × 10^6^/ml were infected with passage 1 of virus. The pellet of the infected cells were collected after 52 hours and lysed in lysis buffer (Tris 50 mM pH 8, NaCl 150 mM, Glycerol 10%, EGTA 1 mM, DTT 4 mM, SigmaFast Protease inhibitor, PMSF 0.1 mM, Benzonase 10 unit, NP-40 0.2%). Clear lysates were incubated with IgG Sepharose 6 Fast-Flow (GE Healthcare) for four hours at 4 °C. The beads were collected and washed with 100 ml lysis buffer (with NP-40 0.02%), then 100 ml high salt lysis buffer (250 mM NaCl) and finally 50 ml of low salt TEV buffer (Tris-HCl pH 8 50 mM, NaCl 50 mM, 10% glycerol). Eventually the beads were incubated with TEV protease in the TEV buffer for 2 hours at 16° C.

In the case of SNAP-BicDR1^FL^-SpyCatcher, the TEV-cut eluent protein wes treated with SNAP-Surface Alexa Fluor® 647. The proteins were further purified by Superose 6 Increase 10/300 GL column (Cytiva) using TEV buffer for equilibration. The fractions collected, concentrated and stored in the same buffer as for gel filtration with 0.5 mM DTT and 10% glycerol.

#### HOOK3

Human Hook3 (1-552) with ZZ-TEV-Halo tag at its N-terminus was received as a gift from Reck-Peterson Lab (UCSD) in pET28a vector. Proteins were expressed in BL21(DE3) *E*.coli strain in autoinduction media (Terrific Broth Base including Trace elements - Formedium) at 25 °C. The cell pellet were lysed at HEPES pH 7.4, 50 mM, NaCl 150 mM, Glycerol 10%, EGTA 1 mM, BME 20 mM, SigmaFast protease inhibitor, PMSF 0.1 mM, Benzonase 10 unit and Lysozyme 5 mgr. Cleared lysate was incubated with washed IgG Sepharose 6 Fast-Flow (GE Healthcare) for 3 hours at 4 °C. The beads were collected and washed in tandem with lysis buffer, high salt lysis buffer (300 mM NaCl), lysis buffer and eventually low salt TEV buffer (Tris.HCl, pH 8 25 mM, NaCl 50 mM, glycerol 5%, EGTA 0.5 mM, DTT 0.5 mM). The proteins released from the IgG beads after overnight incubation with TEV protease in the low salt TEV buffer. For fluorescently labeled Hook3, 300 µl of eluant (at 1.5 mg/ml) was treated with 6.5 µl of 1 mM HaloTag Alexa Fluor 488 Ligand (Promega PAG1001). Both labeled and unlabeled proteins were subject to size exclusion chromatography for further purification using Superose 6 Increase 10/300 GL column (Cytiva) equilibrated in Tris.HCl pH 8 50 mM and 50 mM NaCl. The fractions collected, concentrated and stored at −80 in Tris.HCl pH 8 50 mM and 50 mM NaCl with 6% glycerol.

#### NINL

NINL (1-702) containing an amino-terminal HaloTag was expressed in BL-21[DE3] cells (New England Biolabs), which were then grown until OD 0.4-0.6 and induced with 0.1 mM IPTG for 16 hr at 16°C. Frozen cell pellets from a 2 L culture were resuspended in 60 lL of activator-lysis buffer (30 mM HEPES [pH 7.4], 50 mM potassium acetate, 2 mM magnesium acetate, 1 mM EGTA, 1 mM DTT, 0.5 mM Pefabloc, 10% (v/v) glycerol) supplemented with 1 cOmplete EDTA-free protease inhibitor cocktail tablet (Roche) per 50 ml and 1 mg/ml lysozyme. The resuspension was incubated on ice for 30 min and lysed by sonication. The lysate was clarified by centrifuging at 66,000 x g for 30 minutes in Type 70 Ti rotor (Beckman). The clarified supernatant was incubated with 2 ml of IgG Sepharose 6 Fast Flow beads (Cytiva) for 2 hr on a roller. The beads were transferred to a gravity flow column, washed with 100 ml of activator-lysis buffer supplemented with 150 mM potassium acetate followed by 50 ml of cleavage buffer (50 mM Tris–HCl [pH 8.0], 150 mM potassium acetate, 2 mM magnesium acetate, 1 mM EGTA, 1 mM DTT, 0.5 mM Pefabloc, 10% (v/v) glycerol). The beads were then resuspended and incubated in 15 ml of cleavage buffer supplemented with 0.2 mg/ml TEV protease overnight on a roller. The supernatant containing cleaved proteins was concentrated using a 30K MWCO concentrator (EMD Millipore) to 1 ml, filtered by centrifuging with Ultrafree-MC VV filter (EMD Millipore) in a tabletop centrifuge, diluted to 2 ml in Buffer A (30 mM HEPES [pH 7.4], 50 mM potassium acetate, 2 mM magnesium acetate, 1 mM EGTA, 10% (v/v) glycerol, and 1 mM DTT) and injected onto a MonoQ 5/50 GL column (Cytiva) at 1 ml/min. The column was pre-washed with 10 CV of Buffer A, 10 CV of Buffer B (30 mM HEPES [pH 7.4], 1 M potassium acetate, 2 mM magnesium acetate, 1 mM EGTA, 10% (v/v) glycerol, and 1 mM DTT) and again with 10 CV of Buffer A at 1 ml/min. To elute, a linear gradient was run over 26 CV from 0-100% Buffer B. The peak fractions containing Halo-tagged activating adaptors were collected and concentrated using a 30K MWCO concentrator (EMD Millipore) to 0.2 ml and diluted to 0.5 ml in GF150 buffer (25 mM HEPES [pH7.4], 150 mM potassium chloride, 1 mM magnesium chloride, 1 mM DTT) and further purified via size exclusion chromatography on a Superose 6 Increase 10/300 GL column (Cytiva) with GF150 buffer at 0.5 ml/min. The peak fractions were collected, buffer was exchanged into a GF150 buffer supplemented with 10% glycerol, concentrated to 0.2-1 mg/mL using a 30K MWCO concentrator (EMD Millipore), and flash frozen in liquid nitrogen.

For fluorescent labeling of the HaloTag, the purified protein was incubated with 50 µM Halo-Alexa488 (Promega) for 10 minutes at room temperature. Free dye was removed using a micro bio spin chromatography column (BioRad) equilibrated in GF150 buffer supplemented with 10% glycerol. The labeled protein was flash-frozen in liquid nitrogen.

### Preparation of fluorescent HIV-1 cores and capsids

#### HIV-1 core purification

GFP-labeled HIV-1 cores were purified from HIV-1 particles generated from the Env-defective proviral clone (R9ΔE) and a plasmid expressing the GFP-Vpr fusion protein^6^. Viruses were produced by transfection of 4 x 10^6^ 293T cells seeded in 100 mm dishes with 10 mg of R9ΔE plasmid and 1 mg of GFP-Vpr plasmid. HIV-1 cores were purified from virions concentrated from 30 ml of culture supernatant as described previously^57^. Aliquots of purified cores were snap frozen and stored at −80°C. Samples were thawed on ice prior to use in motility assays.

#### Recombinant CA purification

HIV-1 CA(A14C/E45C), CA(A92E/A204C) and CA(A92E/A204C/R18G) in pET21a plasmid and CA(K156C) in pET11a plasmids were expressed and purified from the *E.* coli BL21(DE3) strain. The transformed bacteria grow to OD_600_ ≈ 2 in hyper broth media before induction by IPTG 0.5 mM. The cells were collected after overnight shaking at 23 °C. Purification steps were followed as explained before^58^. Lysis buffer included 50 mM Tris-HCl at pH 8, NaCl 50mM, Lysozyme 2 mg/50ml, benzonase 1.5 µl/50ml, 10 mM TCEP, 10 mM DTT and SigmaFast protease^TM^ inhibitors. Solid Ammonium Sulfate (Sigma #A4418) was added slowly to clear lysate to reach 27% saturated Ammonium Sulfate. After 30 minutes of stirring, the pellet of precipitated proteins was collected at 40,000 x g, then resuspended in HEPES 25 mM pH 7.2 and TCEP 15 mM. After overnight dialysis in the same buffer, the solution was centrifuged at 70,000 x g and the supernatant of solubilized proteins was injected onto a HiTrap SP HP cation exchange column (Cytiva). The loaded column was washed with no salt buffer and the CA protein was eluted from the column gradiently with over 15 column-volume of HEPES 25 mM, 1000 mM NaCl. The eluted proteins were dialyzed in the low salt buffer overnight. The final protein sample was concentrated to 10-20 mg/ml and evaluated by SDS-PAGE.

#### CA assembly and fluorescence labeling

Cross-linked CA(A14C/E45C) tube-shaped constructs were formed by three tandem overnight dialysis of CA protein (5-10 mg/ml) as described before^59^: 1) assembling buffer: Tris 25 mM pH 8.0, NaCl 1 M, DTT 30mM. 2) the same as assembling buffer omitting DTT, 3) low salt buffer: Tris 25 mM pH 8.0, NaCl 50 mM. HIV-1 CA cones were formed by dialysis of CA(A92E/A204C) in the assembly buffer (IP6 5 mM, MES 30 mM, pH 6; NaCl 30 mM, and BME 30 mM). The disulfide bonds at the interface of the hexameric units were formed by letting another overnight dialysis in the absence of BME. Eventually, the formed cones were dialyzed in MES 30 mM, pH 6, and NaCl 30 mM to remove the extra IP6. Capped tube-shaped CA(A92E/A204C) and CA(A92E/A204C/R18G) were formed in the same manner as HIV-1 CA cones, just IP6 was replaced by NaCl 1M.

The method by Lau *et al*^60^ was followed to create fluorescent HIV-1 capsid. In summary, CA(K158C) was assembled into tubes in the presence of a labeling buffer (Tris 50 mM, pH 7.4, 2.5 M NaCl, and 0.3 mM TCEP). Then the tube was incubated for 2 minutes with a 2X excess molar ratio of fluorescent dye (DyLight™ 488 Maleimide) at room temperature in the dark. Extra dye was quenched with 25 mM βME (final), and the labeled CA tubes were pelleted and resuspended in the fresh labeling buffer. After several steps of pelleting and resuspending to remove the unreacted dye, the tubes were disassembled in the no-salt labeling buffer. Fluorescently labeled CA (K158C) was aliquoted and stored at −80 °C.

To label the HIV-1 CA tubes or cores, labeled CA (K158C) was mixed with unlabeled CA at 1:40 or 1:80 molar ratio before assembling. After the final dialysis, the tubes or cones were centrifuged at 17,000 x g for 10 minutes to remove the free CA (K158C). The pellet was resuspended in HEPES 50 mM, pH 7.2, NaCl 150 mM. Labeled CA-tubes were stored at 4 °C in the dark. For CA cones, the resuspended cones were centrifuged at 4,000 x g for 5 min. The supernatant included labeled “soluble” cones and was stored at 4 °C. The integrity of tubes and cones was evaluated by negative staining. The concentration of tube and cone (as assembled particles) was calculated based on the final CA contraction and the length of tubes/cones as measured by ImageJ ^61^.

### TIRF imaging

#### HIV-1 microtubule recruitment and motility assay

Single-molecule assy and microtubule-binding assays were performed in flow chambers assembled as described previously^62^. Biotin-PEG-functionalized coverslips were made in the lab following described procedures^63^ with some modifications. In summary, coverslips (1.5 refractive index) were sonicated in ethanol 200 proof at 40 °C for 10 minutes. After washing with milliQ water, the coverslips were sonicated in KOH 200 mM at 40 °C for 20 min, followed by washing with milliQ and another round of sonication with ethanol (Fisher, lot# A995-4, HPLC grade). The incubation solution was prepared by mixing methanol (Fisher, lot:A452-4), Acetic Acid (Fisher, A35-500), and aminosilane (lot: A01W32000610) in 20:5:1 volumetric ratio. Coverslips from the last step were incubated overnight in the incubation solution in the dark and dry place. Then, coverslips were washed with water and ethanol and dried out before treating with PEG-biotin solution. PEG solution included 0.25 g/ml mPEG-Succinimidyl Valerate, MW 2000 (Lysanbio), 0.02 g/ml Biotin-PEG-SVA, MW 5000 (Lysanbio) in 0.84% w/v Sodium bicarbonate solution. 60 µl of PEG solution was added to a clean ethanol-washed slide, and one of the above coverslips was laid down on that 60 µl solution and incubated for 2-3 hours in the dark. Then the coverslips and slides were washed with milliQ water and kept in the pair in a vacuum for future use. Biotin-PEG-functionalized coverslips were stable for at least 1.5 months after preparation.

Microtubules with ∼5% biotin–tubulin (Cytoskeleton inc.) and ∼5% Alexa488-labeled or Alexa647-labeled fluorescent tubulin (Cytoskeleton inc.) were prepared as described^64^ and stabilized with final 20µM taxol (Cytoskeleton inc.). The flow chamber was incubated sequentially with 1 mg/ml streptavidin and then with diluted taxol-stabilized microtubules (final tubulin at 0.025 mg/ml) with 5 minute incubation time. After each step, the chamber was washed twice with the BRB80 buffer (K-PIPES 80 mM, EGTA 1 mM, MgCl2 2 mM, pH 6.8, supplemented with 20 μM taxol).

Single-molecule TIRF microscopy was performed using Olympus inverted microscope at 60X magnification with no further magnifier. The microscope was equipped with Hamamatsu CCD digital camera at 16µm pixel size. Olympus cellTIRF-4Line system provided four laser channels with independent beam paths and depth. Semrock single-band Brightline filters were used for the excitation path filter, and 405/488/561/647 nm Laser Quad Dichromic cube (TRF89902 - ET, Chroma) was the integral emission filter. Olympus CellSens Dimension software controlled the image and movie acquisition, and the z-focus was controlled manually. In all experiments, laser power was set to 15%-25% for all laser beams, which measured 3.5-9.2 mW for 488 nm laser, 2.4-5.8 mW for 561 nm laser, and 4.2-12.9 mW for 647 nm laser.

All imaging was acquired in the DLB buffer (HEPES 25mM, pH 7.4, Mg-Acetate 2 mM, Glycerol 10%, EGTA 1 mM) and 20 µM taxol. Mg-ATP was kept at 1 mM unless otherwise noted. For consistency, no reducing agent was used in the imaging solution. The oxygen scavenging system was based on gloxy combination of glucose oxidase and catalase enzymes^65^.

The complexes were assembled by mixing HIV-1 cores (GFP-Vpr) or fluorescently labeled HIV-1 cone-shaped/tube-shaped capsids with dynein for 10 minutes on ice, then dynactin and activating adapter added and incubated for an additional 10 minutes. Core:activator:dynactin:dynein were mixed in 1:200:500:1000 ratio. The final salt concentration in all experiments was kept at 30-35 mM.

HIV-1 (core or capsid) and microtubules were imaged with 500 ms intervals and 50 ms exposure for each fluorescent channel. All single molecule assays were recorded for 4 minutes unless otherwise mentioned. Velocities, bindings and processivity were calculated from kymographs using Fiji/ImageJ^66^ as described before^67^. The total number of bindings per microtubule (at each field of view for each replicate of the experiment) was divided per imaging time (in minutes), length of the microtubule (in µm), and the concentration of HIV-1 core or capsid (full complex, in nM) to calculate the binding rate without any further normalization. Binding and velocity data were collected from at least 20 (in average 25) individual, fully extended microtubules.

#### Competition experiments

To figure out the binding site(s) of dynein on HIV-1 capsid, motility assay was performed in presence of inositol six-phosphate, IP6 (Phytic acid sodium salt hydrate, Sigma-Aldrich #P8810) and PF74 (PF-3450074, MedKoo Biosciences #555391). IP6 is known to coordinate rings of electropositive charge in the central pores of HIV-1 hexamers that interact with K25 and R18 rings at the pore. PF74 is a small molecule that binds to FG binding pocket. Cone-shaped HIV-1 capsid was assembled with IP6 and fluorescently labeled as described in earlier sections. IP6 eventually was removed by dialysis, pelletting and resuspension. The motility assay was set up with Dynein (10 nM) dynactin and BicDR1 as described before. IP6 (made at 500 mM stock solution and pH adjusted to 6) was added as the last component to the imaging mixture at 10 or 100 μM final concentration and incubated on ice for 10 min before recording HIV-1 capsid motility. PF-74 stock solution was made freshly in DMSO. PF74 was also added at final 1 or 10 μM to the imaging solution as the last component and incubated on ice for 10 min before imaging. Velocities, bindings and processivity were calculated and compared in presence of each putative competitor.

### Encapsulin viral-like model system

#### Engineered encapsulin: design and purification

A synthetic two-gene operon for Encapsulin-SpyTag expression was designed containing mNeonGreen, N-terminally fused to the native IMEF cargo protein from Quasibacillus thermotolerans (UniprotID: A0A0F5HNH9), separated by a GGSGGS linker, and the Q. thermotolerans encapsulin (UniprotID: A0A0F5HPP7) with a C-terminal SpyTag (SAHIVMVDAYKPTK). The synthetic operon was codon optimized for E. coli overexpression and ordered as a gBlock gene fragment from Integrated DNA Technologies (IDT). The gBlock was designed to contain a 20 bps overlap with the multiple cloning site 2 (MCS2) of the pETDuet-1 expression vector. The expression plasmid was assembled via Gibson Assembly by mixing pETDuet-1 vector digested with NdeI and PacI with the gBlock in addition to Gibson Assembly Mix (NEB #E2611L) and incubating at 50°C for 15 min. Assembled plasmid was inserted into electrocompetent E. coli DH10B cells by transformation via electroporation. The plasmid was confirmed via Sanger sequencing (Eurofins).

For expression of the Encapsulin-SpyTag protein, 500 ml of terrific broth (TB) in a 2 L baffled flask containing 100 µg/ml of ampicillin was inoculated with a 10 ml overnight culture of *E.coli* BL21 (DE3) containing the Encapsulin-SpyTag expression plasmid. The culture was grown with shaking at 200 rpm at 37°C until it reached an OD600 of 0.5. Protein expression was then induced by the addition of 0.2 mM IPTG and the culture was then moved to 30°C and grown with shaking at 200 rpm for 22 hours. Cells were then harvested by centrifugation and the cell pellet was immediately lysed by sonication in a lysis buffer consisting of 150 mM NaCl, 20 mM Tris pH 8.0, 1 mM MgCl_2_, 1 mM CaCl_2_, 1x SigmaFast EDTA-free protease inhibitor cocktail, 0.5 mg/ml lysozyme, and 10 µg/ml of DNAse I. Sonication was performed using a Model 120 Sonic Dismembrator (Fisher Scientific, Inc.) at 48% amplitude for 5 minutes total (10 seconds on, 20 seconds off) on ice. Cell debris was removed from the lysate by centrifugation at 10,000 x g for 10 minutes at 4°C. The clarified supernatant was then incubated on ice with gentle rocking for 30 minutes. Solid PEG-8000 (10% w/v final concentration) and 0.6 M NaCl were added to the supernatant, followed by incubation with gentle rocking at 4°C for 45 minutes. The precipitate was pelleted by centrifugation at 8,000 x g for 15 minutes at 4°C, resuspended in 5 ml of 150 mM NaCl, 20 mM Tris pH 8.0, and filtered with a 0.2 µm syringe filter. The filtered protein sample was then subjected to size exclusion chromatography on a HiPrep 16/60 Sephacryl S-500 size exclusion column (GE Health Sciences) using a running buffer consisting of 150 mM NaCl, 20 mM Tris pH 8.0. Encapsulin-containing fractions were pooled, concentrated, and desalted into 20 mM Tris pH 8.0 using an Amicon Ultra-15 centrifugal filter (EMD Millipore) with a 100 kDa molecular weight cutoff, and further purified by ion-exchange chromatography using a HiPrep DEAE FF 16/10 ion exchange column (GE Health Sciences). After loading, the column was washed with 100 ml of 20 mM Tris pH 8.0 followed by a 200 ml linear gradient to 1 M NaCl, 20 mM Tris pH 8.0. Encapsulin-containing fractions were again concentrated and exchanged into 150 mM NaCl, 20 mM Tris pH 8.0 using an Amicon Ultra-15 centrifugal filter with a 100 kDa molecular weight cutoff and subjected to a final round of size exclusion chromatography using a Superose 6 Increase 10/300 GL size exclusion column (GE Health Sciences) with 150 mM NaCl, 20 mM Tris pH 8.0 as the running buffer. All chromatography steps were performed using an ӒKTA Pure FPLC system (GE Health Sciences). Encapsulin-containing fractions were then pooled, concentrated to 3.3 mg/mL according to absorbance at 280 nm, and immediately drop frozen in 25-30 µl aliquots in liquid nitrogen. The frozen protein was then stored at −80°C until needed.

#### Encapsulin-BicDR1: motility data

Purified BicDR1^FL^-SpyCatcher was incubated with Encapsulin-SpyTag. The incubation buffer includes 50 mM Tris pH 8.0, 50 mM NaCl, 1 mM EDTA, 9% glycerol, 0.5 mM DTT at the final volume of 21.5 µl. The Encapsulin-SpyTag concentrations were kept constant at 15.5 nM while the concentration of BicDR1^FL^-SpyCatcher adjusted to keep the ratio of 1, 2.5, 10, 25 and 100 BicDR1 (as a dimer) per Encapsulin. The solutions were incubated at 4 °C for at least 24 hours. To evaluate the yield of the reaction and to remove the unreacted BicDR1, the complexes of Encapsulin-BicDR1 (∼9-25 MDa) were diluted and subjected to ultrafiltration using 1 MDa MWCO VIVASPIN. The formation of Encapsulin-SpyTag-BicDR1^FL^-SpyCatcher complexes was analyzed and further quantified by SDS-PAGE. We confirmed the formation of a new band related to Encapsulin-SpyTag-BicDR1^FL^-SpyCatcher even at the lowest concentration of BicDR1. Based on the ultrafiltration and SDS-PAGE, all added BicDR1-SpyCatcher was incorporated into Encapsulin-spyTag.

TIRF experiments of Encapsulin-SpyTag-BicDR1^FL^-SpyCatcher with Enc:BicDR1(dimer) ratio of 1:1, 1:10, 1:100 were conducted using the same buffer condition as was used for HIV-1 (DLB buffer, with 30 mM salt and 1 mM ATP, 20 µM taxol, and gloxy). Dynein and dynactin were kept constant at the saturation concentration, while the concentration of Encapsulin-SpyTag-BicDR1^FL^-SpyCatcher construct was adjusted according to the number of BicDR1 per encapsulin. For example, we calculated how much of the Enc:BicDR1 construct at each ratio needs to be added to keep the final total BicDR1 in the imaging solution at 0.5 nM. In this way, the molarity and overall ratio of dynein, dynactin and BicDR1 remained exactly the same in all single molecule assays, while the concentration of encapsulin itself (the monitored single particles) in the imaging solution was 0.5 nM, 0.05 nM and 0.005 nM for 1:1, 1:10, 1:100 Enc:BicDR1 experiments respectively.

Taxol-stabilized microtubules (labeled with Alexa-647 and 5% biotin-tubulin) were immobilized on the PEG-Biotin treated coverslip as described before in this manuscript. Imagining solution including encapsulin-BicDR1 was introduced into the chamber with immobilized microtubules and immediately imaged by TIRF microscope. A 488 nm laser was used for the excitation in the encapsulin channel (mNeonGreen). All the movies were recorded for 4 minutes with 500 ms interval and 50 ms exposure for 488 nm and 647 nm excitation. The velocity and binding rate were calculated from the extracted kymographs using Fiji/ImageJ^66^.

#### Standard curve creation by encapsulin-BicDR1 and quantification of dyneins per HIV-1 core

Encapsulin-SpyTag-SNAP-BicDR1^FL^-SpyCatcher complex was used to generate the standard curve to calculate the number of dyneins bound to the HIV-1 core. The standard curve is supposed to plot the fluorescent intensity per encapsulin when a different number of fluorescently labeled BicDR1 is bound to encapsulin. To do so, purified SNAP-BicDR1^FL^-SpyCatcher was labeled with SNAP-Surface Alexa Fluor®647 at almost 100% labeling yield. To generate encapsulin-SpyTag-SNAP^Alexa647^–BicDR1^FL^-SpyCatcher the same incubation conditions, preparation and evaluation steps as described for BicDR1^FL^-SpyCatcher were followed. Enc:SNAP^Alexa647^–BicDR1 at 1:1, 1:2.5, 1:10, 1:25 and 1:100 ratio were diluted at least 200 times in the imaging buffer (DLB with 33 mM salt, 1 mM ATP and gloxy system) to make a layer of separated single particles on the untreated cover slip. Then several images were immediately taken at 50 ms exposure from different spots on the coverslip at 647 nm excitation. Images were processed with Fiji/ImageJ software. An ROI was created and applied to all the images in order to measure the fluorescence intensities of single particles at the region away from the edge of the image. The ComDet^68^ plugin installed in Fiji was used to detect spots and analyze colocalization. The ComDet parameters for colocalization analysis including distance between colocalized spots (4 pixel), approximate particle size (4 pix) and intensity threshold (2) were applied and large particles were segmented. The fluorescence intensity (a.u.) at 647 nm were calculated and collected only from the particles co-localized with 488 nm fluorescence intensity (encapsulin) and then normalized by subtracting the background fluorescence intensity at 647 nm for each image. A histogram plot was generated from the normalized intensity data for each of Enc:SNAP^Alexa647^–BicDR1, and mean values were determined by fitting a gaussian curve to histogram data. By plotting these mean values against the number of SNAP^Alexa647^–BicDR1 per encapsulin the standard curve was formed.

WT dynein was also fluorescently labeled with SNAP-Surface Alexa Fluor® 647 via the SNAP-tag at the N-terminus of DHC with 93% labeling efficiency. Then dynein at 10 nM was incubated with 0.05 nM HIV-1 cores (GFP-vpr) in the buffer similar to TIRF assay (DLB with 30 mM salt, 1 mM ATP, gloxy system). The mixture was incubated on ice for 20 min, then diluted 20 times in the same incubation buffer before infusing the solution into a chamber of untreated coverslip. Imaging at 647 nm was performed following exactly the same conditions and microscope settings as used for Encapsulin-SpyTag-SNAP^Alexa647^–BicDR1^FL^-SpyCatcher. Images were processed by applying the same ROI and ComDet parameters as used for Encapsulin-SpyTag-SNAP^Alexa647^–BicDR1^FL^-SpyCatcher. Histogram graphs from the normalized fluorescence intensity at 647 nm for single particles co-localized with HIV-1 core (at 488 nm excitation) were generated and the mean value of this fluorescence intensity was calculated from the fitted Gaussian curve to the histogram graph. The number of dyneins per HIV-core was calculated by using the trendline equation of Encapsulin-SpyTag-Snap^Alexa647^–BicDR1^FL^-SpyCatcher standard curve and applying the labeling efficiency of dynein.

#### *In-vitro* co-sedimentation assays

Cross-linked CA(A14C/E45C) was prepared as described in the previous section. Purified dynein, dynein tail domain, dynein motor domain were centrifuged at 17,000 x g, 4 °C for 30-60 min before co-pelleting. Each protein was added to CA tubes (equal to ∼30 µM CA hexamer) in 40 µl final volume. DLB with 0.1 mM ATP and the final salt of 50 mM was used as an incubating buffer. The mixtures were incubated for 60 min at 4 °C on the roller. Subsequently, 10 μl aliquots were withdrawn and labeled as total. The remaining was centrifuges at 4 °C for 7 min at 40,000 x g. The pellet resuspended in 30 µl of loading buffer + BME. 10 µl of total, supernatant, and pellet samples were analyzed by SDS-PAGE.

#### Electron Microscopy

All negative stain imaging in this paper was performed using a Morgagni 268(D) transmission electron microscope S3 microscope (FEI Co.). The microscope is equipped with a tungsten filament operated at 100 Kv high tension, and a Gatan Orius SC200 CCD Detector with a physical pixel size of 7.4 µm. Images well acquired at nominal magnification of 22,000x (2.1 Å/pixel) at room temperature. The samples were fixed using conventional negative staining procedures with 0.075% uranyl formate on glow-discharged grids with Formvar carbon film (Electron Microscopy Sciences, FCF-400-Cu-50).

### Cell-based HIV-1 infection assays

#### siRNA knockdowns

siRNA targeting BicDR1 (Ambion siRNA #149588), Hook3 (Ambion siRNA #s228364, or negative control siRNA (Negative Control No. 1, Thermofisher #4390843) was purchased from Thermofisher Scientific. siRNAs were transfected twice into cells 24 hours apart using Lipofectamine 3000 (Invitrogen) as per the manufacturer’s protocol. In brief, 300pmol of siRNA was added to OptiMEM media containing 7.5uL of Lipofectamine 3000 and incubated for 15min at room temperature. Following incubation, the siRNA mixture was added dropwise to cells and allowed to incubate for 12 hours in media without antibiotics before replacing media. Cells were allowed to rest for 8 hours before the second transfection was performed using the same protocol. Following the conclusion of the second transfection, cells were collected for confirmation of knockdown via qPCR or western blot as well as subsequent experiments.

To confirm siRNA knockdown western blot or qPCR analysis was performed. For western blot, siRNA-treated cells were lysed 12-hours following the second siRNA transfection. Western blot was performed using the following antibodies: mouse Anti-Hook3 (sc-398924). For qPCR, RNA was isolated and purified from cells using the NucleoSpin RNA extraction kit (Macherey-Nagel). Following RNA isolation, cDNA conversion was performed using the GoScript Reverse Transcriptase Kit (Promega). qPCR was performed using separate sets of primers amplifying various regions of each gene. The primer sets were as follows:

BICDR1 #1 Fwd, 5’-GAG CTG GAG AGT GAT GTG AAG-3’, BICDR1 #1 Rev, 5’-CCT GCT GAG CTG ATC CAA TAG-3’; BICDR1 #2 Fwd, 5’-CAT CAA CCA ACC AGC ACA TTA TC-3’, BICDR1 #2 Rev, 5’-TCC TCT AAA GTA GCG CTG AGA-3’; BICDR1 #3 Fwd, 5’-CAA GTC ATC TGC AGG CTT TA-3’, BICDR1 #3 Rev, 5’-GTT TGG CTA GGC CAA GAA ATT AT-3’; BICDR1 #4 Fwd, 5’-CGA GCA CTT AGA GCA AGA GAA A-3’, BICDR1 #4 Rev, 5’-GTA GCT GCT TCA CAT CAC TCT C-3’; BICDR1 #5 Fwd, 5’-ACA TCC CTC CTG TCA GAG AT-3’, BICDR1 #5 Rev, 5’-CGA AGG TGT GAG CAC AGA TAG-3’.

Hook3 #1 Fwd, 5’-GAT CAG TAG TGA GTG AGT GTT AAG T-3’, Hook3 #1 Rev, 5’-AGG AGG AAG GAA GGA GAG ATA G-3’, Hook3 #2 Fwd, 5’-GTG GAG TGG ATC ATT GGG ATA AG-3’, Hook3 #2 Rev, 5’-CTA TTG CTT TGG GAA CAT CTG TTT AG-3’, Hook3 #3 Fwd, 5’-AGG AGC AGC ACC AGA AAT AC-3’, Hook3 #3 Rev, 5’-CTT CCA TCT CTC TCT GAG TCT TTG-3’

#### Virus Production and Infection analysis

To generate HIV-1 particles pseudotyped with VSVg, 293T cells seeded in a 10-cm dish at ∼70% confluency was transfected with either 7ug of R7ΔEnv-GFP or NL4.3Luc-mCherry and 3ug of pCMV-VSVg using polyethylenimine (PEI). Media was replaced 12 hours after transfection. Viruses were harvested 48 hours after transfection and filtered through a 0.45-um filter (Milipore). To measure infectivity, equal amounts of NL4.3Luc-mCh or R7-GFP virus, as measured by RT activity, was added to cells and a synchronized infection was performed by spinoculation at 13°C for 2 hours at 1,200 x g. After spinoculation, virus-containing media was removed and replaced with 37°C media. Infectivity was measured 48 hours after infection using either luciferase assay to measure presence of firefly luciferase or flow cytometry to measure the number of GFP-positive cells.

#### Proximity Ligation Assay

A Duolink Pla kit was purchased and assayed per the manufacturer’s protocol. In brief, cells were grown on coverslips and fixed with 3.7% paraformaldehyde at time points following synchronized infection. To determine interaction between BicDR1 and the viral capsid protein p24 cells were permeabilized and blocked using 3% donkey serum followed by incubation with primary antibodies targeting p24 (Santa Cruz sc-69728) and BicDR1 (Atlas Antibodies HPA041309). After primary staining, coverslips were washed and incubated (1 hour at 37°C) with secondary anti-mouse conjugated minus and anti-rabbit conjugated plus Duolink PLA probes. Following incubation coverslips were washed once again and incubated with ligation-ligase solution (30 min at 37°C) followed by subsequent incubation with amplification-polymerase solution (100 min at 37°C) containing Duolink Red reagents. Upon completing the PLA protocol, coverslips were additionally incubated with PBS containing Dapi and phalloidin-488 and subsequently mounted.

Z-stack images were acquired using a DeltaVision wide-field fluorescent microscope (Applied Precision, GE) equipped with a CoolSNAP camera (CoolSNAP HQ; Photometrics) using a 60x objective lens. Excitation light was generated using an illumination module (Applied Precision, GE) and deconvolved using SoftwoRx deconvolution software. All images were imaged using identical acquisition parameters. Following acquisition PLA puncta were quantified using Imaris software.

#### Statistical analyses

All data were analyzed and plotted by Prism 9 or Prism 10. Two-tailed unpaired t-test with Welch’s correction (did not assume equal standard deviation for both populations) was used to calculate p-values for pairwise comparison. For non-normal distribution non-parametric Mann-Whitney t-test was used. In order to conduct multiple comparisons, a normality test (based on D’Agostino-Pearson normality test with 0.05 significance level) was first conducted to determine if the data follows a normal (Gaussian) distribution. If all the datasets followed the normal distribution, statistical analyses were performed using one-way Brown–Forsythe and Welch ANOVA with correction for multiple comparison using the Dunnett T3 test. And the non-parametric test was conducted if one or more of the datasets were not normally distributed. Dunn’s test was used to correct for non-parametric ANOVA. Multiple compassion followed the mean rank of each column with the mean rank of the control column or with every other column. The *p*-values are reported on all graphs with asterisks following the GraphPad style with 95% confidence level: ns = P >0.05, * = P ≤0.05, ** = P ≤0.01, *** = P ≤0.001, **** = P ≤0.0001.

## Notes

### Competing Interest Statement

The authors have declared no competing interest.

### Summary of Updates

We have updated the text and included a new figure for a final model of HIV-1:dynein.

