## Supplemental Figures for "HIV-1 binds dynein directly to hijack microtubule transport machinery"

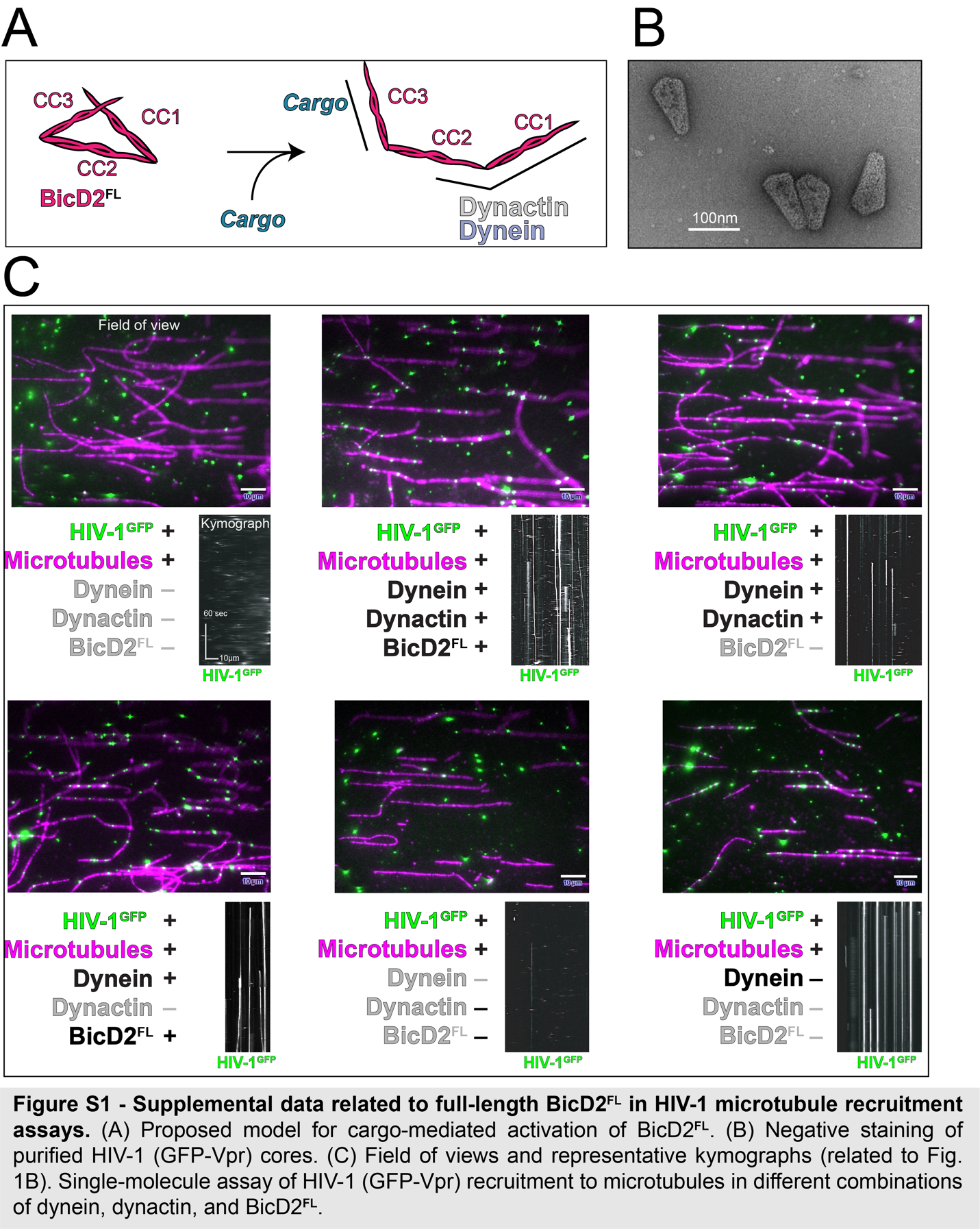

Fig. S1.

**Supplemental data related to BicD2^FL^ with HIV-1 in microtubule recruitment assays.** **(A)** Proposed model for cargo-mediated activation of BicD2^FL^. **(B)** Negative staining of purified HIV-1 (GFP-Vpr) cores. **(C)** Field of views and representative kymographs (related to Fig. 1B). Single-molecule assay of HIV-1 (GFP-Vpr) recruitment to microtubules in different combinations of dynein, dynactin, and BicD2^FL^.

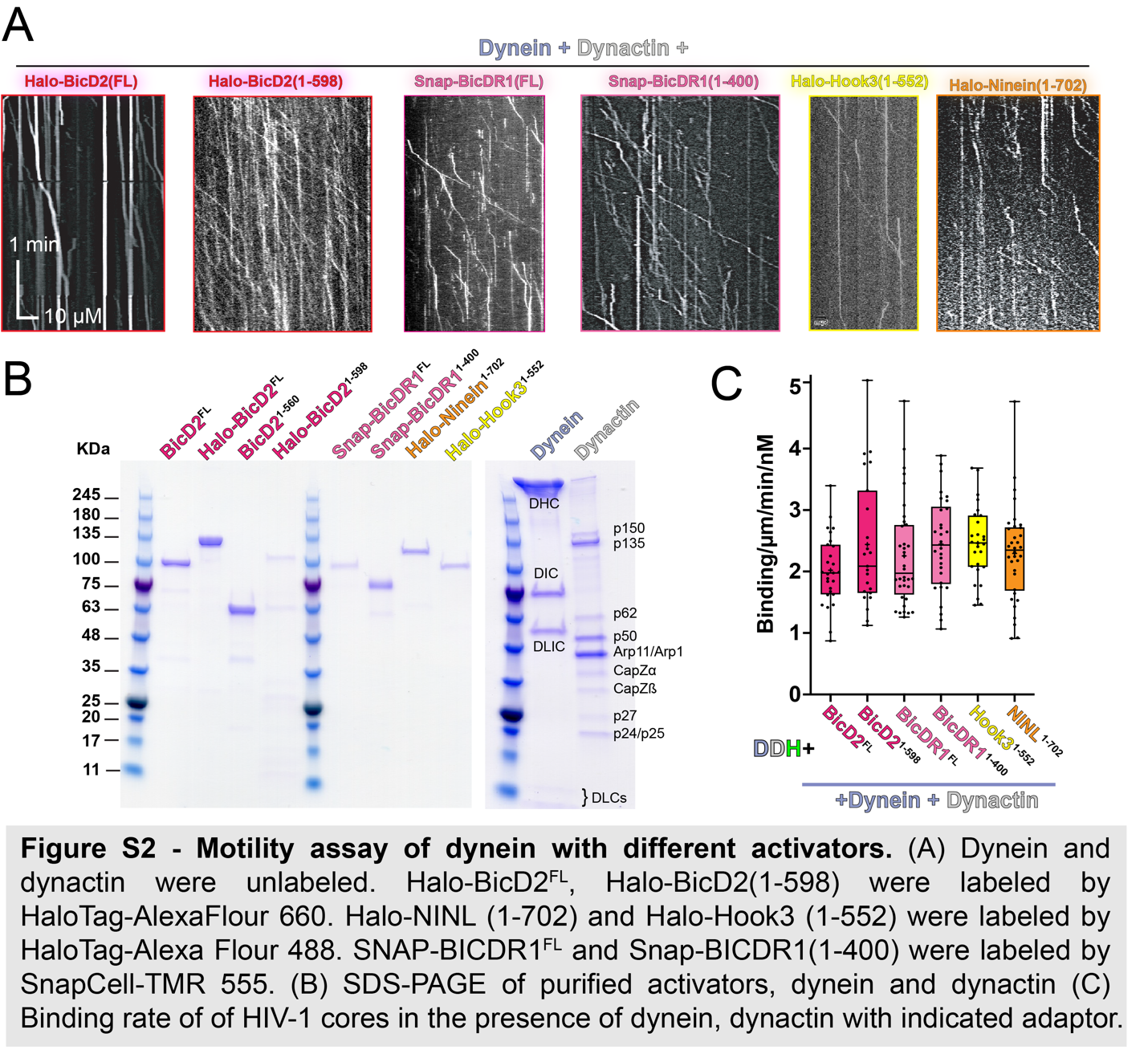

Fig. S2.

**Motility assay of dynein with different activators.** **(A)** Motility of dynein-dynactin with indicated adaptor. Dynein and dynactin were unlabeled. Halo-BicD2^FL^, Halo-BicD2^1-598^ were labeled by Halo-AlexaFlour 660. Halo-NINL^1-702^ and Halo-Hook3^1-552^ were labeled by HaloTag-Alexa Flour 488. SNAP-BicDR1^FL^ and SNAP-BicDR1^1-400^ were labeled by SnapCell-TMR **(B)** SDS-PAGE of purified activators, dynein and dynactin **(C)** Binding rate of HIV-1 cores in the presence of dynein, dynactin with indicated adaptor.

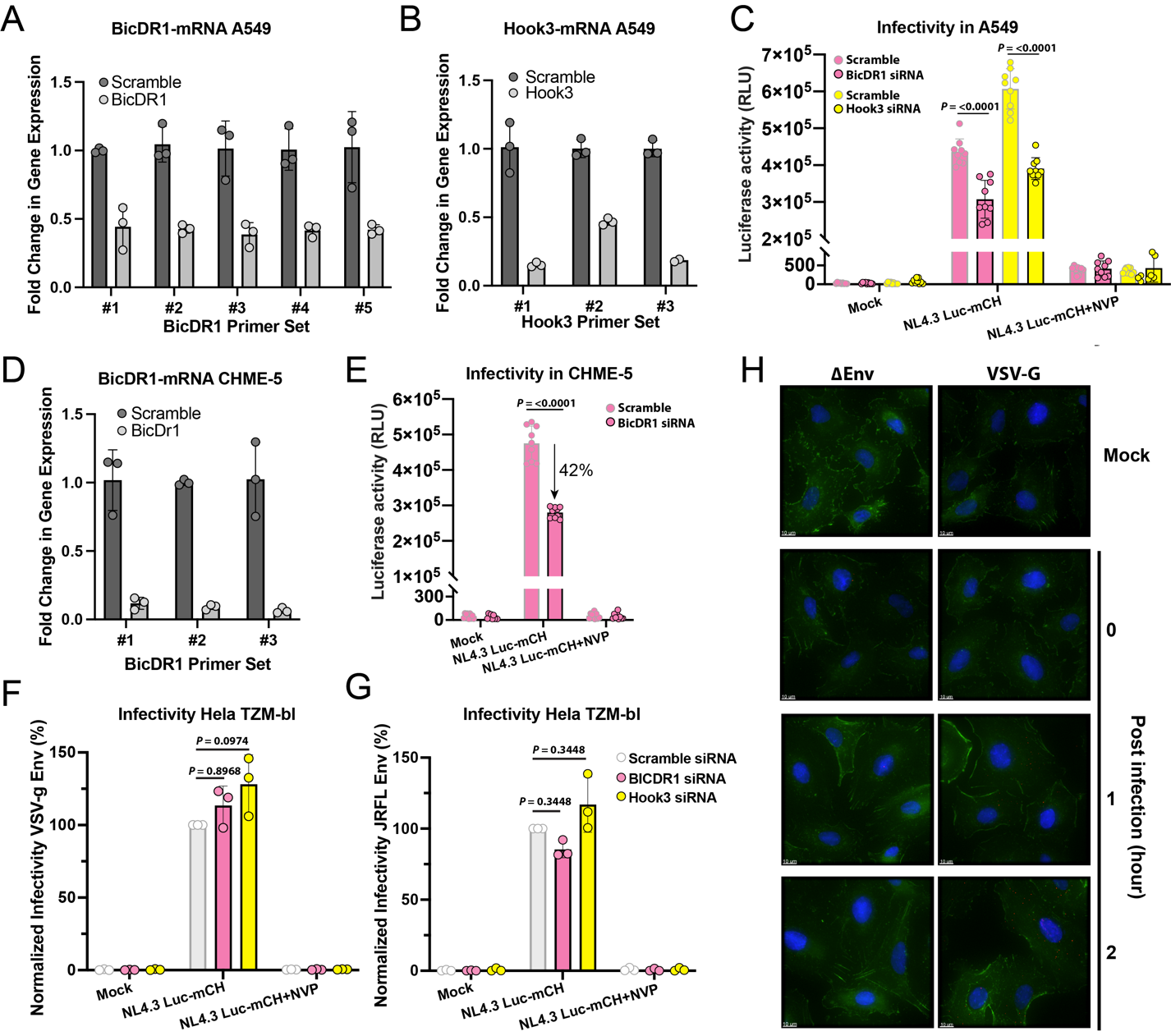

Fig. S3.

**BicDR1 and Hook3 are required for HIV-1 infection.** **(A)** BicDR1 mRNA level in BicDR1 depleted A549 cells as tested by qPCR using 5 different primer sets. **(B)** Hook3 mRNA level in Hook3 depleted A549 cells as tested by qPCR using 3 different primer sets. **(C)** A549 cells lacking BicDR1 or Hook3 were infected with NL4.3 Luc-mCH virus pseudotyped with VSV-G envelope glycoprotein. Cells were harvested 48 hours post infection and luciferase activity was measured. Luciferase activity was also measured post-infection when cells were treated by Reverse Transcriptase inhibitor, Nevirapine (NVP). Data points (±SD) from nine independent measurements are plotted. **(D)** BicDR1 mRNA level in BicDR1 depleted CHME-5 cells as tested by qPCR using 3 different primer sets. **(E)** CHME-5 cell lacking BicDR1 were infected with NL4.3 Luc-mCH virus pseudotyped with VSV-G envelope glycoprotein. Luciferase activity was measured 48 hours post infection. As a control, infection in presence of Reverse Transcriptase inhibitor, Nevirapine (NVP), was also measured. Data points (±SD) from three independent measurements are plotted. **(F and G)** Normalized infectivity (%) in Hela TZM-bl depleted in BicDR1 or Hook3. The cells were infected with NL4.3 Luc-mCH virus either pseudotyped with VSV-G envelope glycoprotein (F) or expressing CCR5 tropic JRFL (G). **(H)** Related to Fig 2G, Proximity Ligation Assay to determine interaction between BICDR1 and the viral capsid protein p24. A549 cell were infected with R7ΔEnv virus pseudotyped with envelope glycoprotein VSV-G. Santa Cruz sc-69728 antibody was used to target p24 and Atlas Antibodies HPA041309 was used to detect BicDR1.

Statistical analyses were performed using non-parametric one-way ANOVA for multiple comparison (F and G) and Mann-Whitney non-parametric t-test for pairwise comparison (C and E).

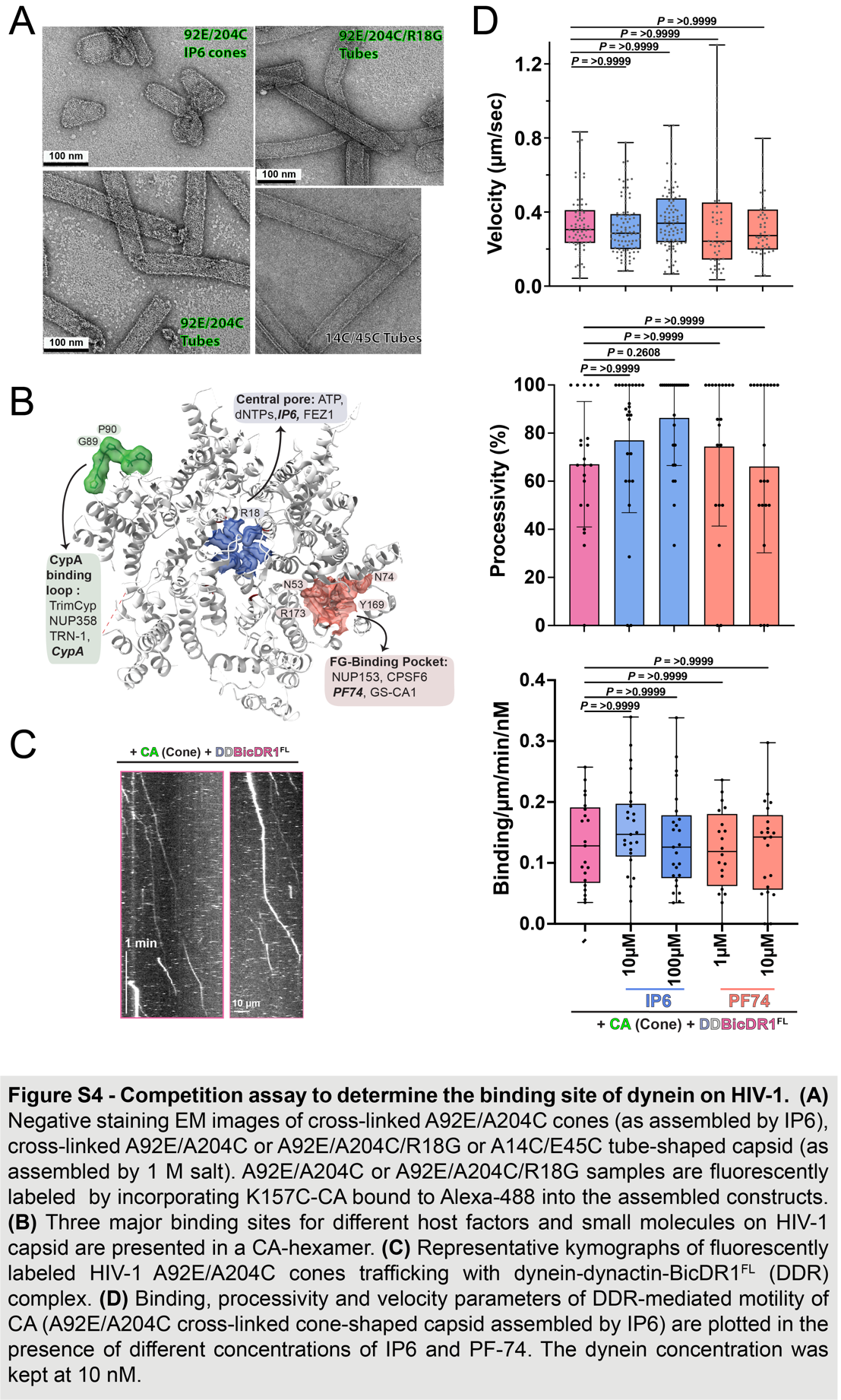

Fig. S4.

**Competition assay to determine the binding site of dynein on HIV-1.** **(A)** Negative staining EM images of cross-linked A92E/A204C cones (as assembled by IP6), cross-linked A92E/A204C or A92E/A204C/R18G or A14C/E45C tube-shaped capsid (as assembled by 1 M salt). A92E/A204C or A92E/A204C/R18G samples are fluorescently labeled by incorporating K157C-CA bound to Alexa-488 into the assembled constructs. **(B)** Three major binding sites for different host factors and small molecules on HIV-1 capsid are presented in a CA-hexamer. **(C)** Representative kymographs of fluorescently labeled HIV-1 A92E/A204C cones trafficking with dynein-dynactin-BicDR1^FL^ (DDR) complex. **(D)** Binding, processivity and velocity parameters of DDR-mediated motility of CA (A92E/A204C cross-linked cone-shaped capsid assembled by IP6) are plotted in the presence of different concentrations of IP6 and PF-74. The dynein concentration was kept at 10 nM.

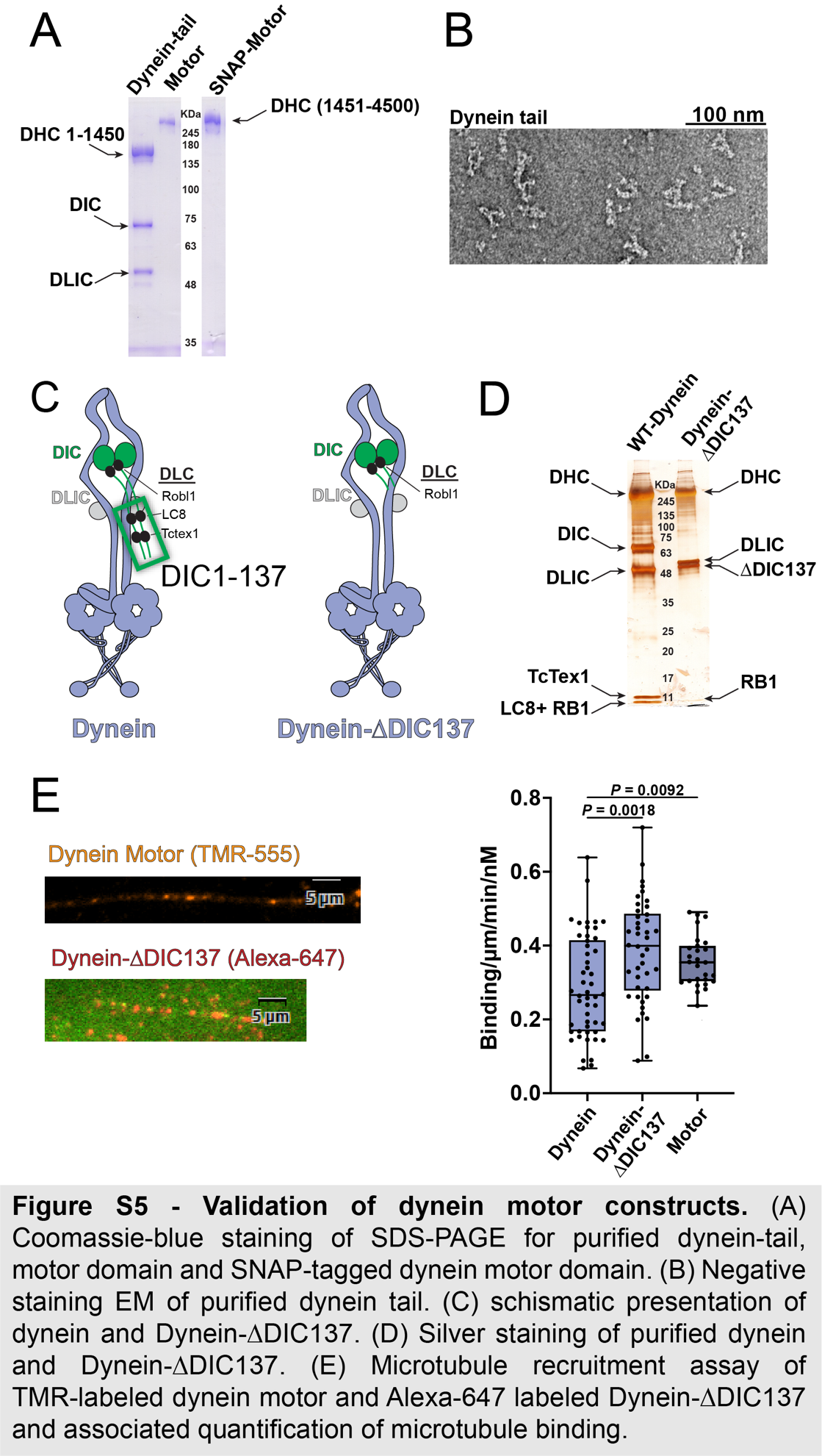

Fig. S5.

**Validation of dynein motor constructs.** **(A)** Coomassie-blue staining of SDS-PAGE for purified dynein-tail, motor domain and SNAP-tagged dynein motor domain. **(B)** Negative staining EM of purified dynein tail. **(C)** Schematic presentation of Dynein and Dynein-∆DIC137. **(D)** Silver staining of purified dynein and Dynein-∆DIC137. **(E)** Microtubule recruitment assay of TMR-labeled dynein motor and Alexa-647 labeled Dynein-∆DIC137 and associated quantification of microtubule binding.

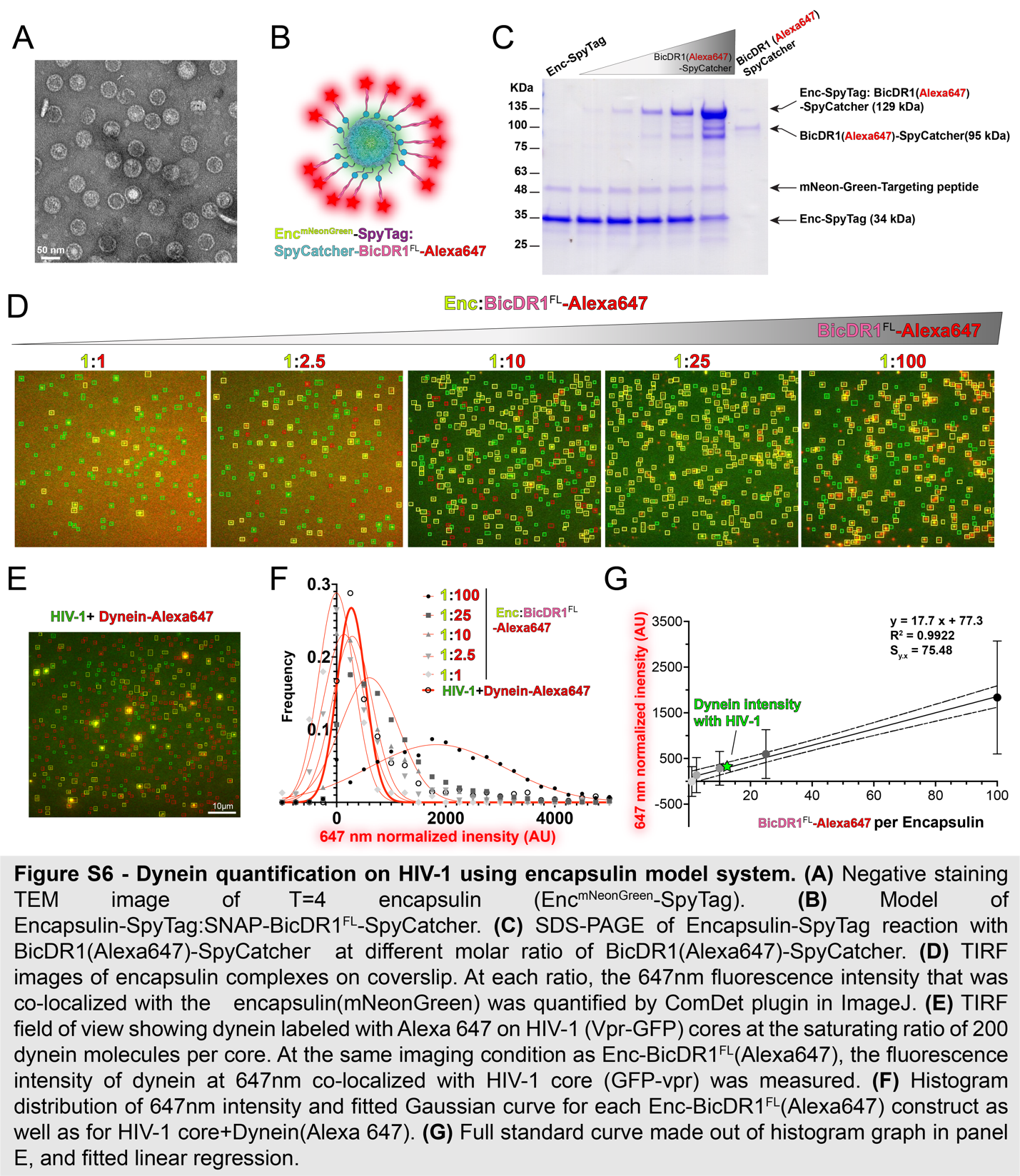

Fig. S6.

**Dynein quantification on HIV-1 using encapsulin model system.** **(A)** Negative staining TEM image of T=4 encapsulin (Enc-mNeonGreen-SpyTag). **(B)** Model of Encapsulin-SpyTag-SNAP ^Alexa647^-BicDR1^FL^-SpyCatcher. **(C)** SDS-PAGE of Encapsulin-SpyTag reaction with SNAP^Alexa647^-BicDR1^FL^-SpyCatcher at different molar ratio of SNAP^Alexa647^-BicDR1^FL^-SpyCatcher. **(D)** TIRF images of encapsulin complexes on coverslip. At each ratio, the 647 nm fluorescence intensity that was co-localized with the encapsulin(mNeonGreen) was quantified by ComDet plugin in Fiji/ImageJ. **(E)** TIRF field of view showing dynein labeled with Alexa 647 on HIV-1 (Vpr-GFP) cores at the saturating ratio of 200 dynein molecules per core. At the same imaging condition as Enc- SNAP^Alexa647^-BicDR1^FL^, the fluorescence intensity of dynein at 647 nm co-localized with HIV-1 core (GFP-vpr) was measured. **(F)** Histogram distribution of 647 nm intensity and fitted Gaussian curve for each Enc- SNAP^Alexa647^-BicDR1^FL^ construct as well as for HIV-1 core + Dynein^Alexa647^. **(G)** Full standard curve made out of histogram graph in panel E, and fitted linear regression.
